# Isl1^+^ Central Amygdala Neurons Coordinate Control of the Jaw and Stomach During Ingestion

**DOI:** 10.64898/2026.08.11.744231

**Authors:** Matthew H. Perkins, Wenfei Han, Leonardo Santana Novaes, Hao Chang, Ivan de Araujo

## Abstract

Central Amygdala neurons expressing Isl1 (CeA^Isl1+^) project to brainstem regions involved in control of the jaw and the stomach, including the parabrachial nucleus (PBN), the nucleus of the tractus solitary (NTS), and the parvocellular reticular nucleus (PCRt). Stimulation of CeA^Isl1+^ cells elicits fictive feeding, particularly biting. Activation of these neurons can rate dependently set the amplitude of bite force and inhibition dramatically reduces bite force. Findings suggest this force generation depends on modulation of a jaw closing reflex involving tooth sensory neurons in the mesencephalic trigeminal nucleus (Me5). Anatomical tracing studies show Me5 neurons receive synaptic input from CeA^Isl1+^ neurons. Patch clamp recordings of Me5 neurons indicate this synapse is mediated by GABA yet depolarizing. Activation of CeA^Isl1+^ neurons is capable of dramatically potentiating the periodontal jaw closing reflex, a reflex whereby Me5 tooth sensory neurons activate jaw closing muscles. In addition to controlling the actions of the jaw, CeA^Isl1+^ neuron stimulation is sufficient to reduce gastric pH. Inhibition experiments show these cells are necessary for lowering gastric pH in mice anticipating a meal. Finally, CeA^Isl1+^ neurons can modulate gastric motility, stimulation transiently suppresses gastric motility, an effect also observed when animals chewed food. Subdiaphragmatic vagotomy eliminated the transient suppression of gastric motility otherwise observed with CeA^Isl1+^ neuron stimulation or food chewing. Taken together, this molecularly and anatomically defined population generates specific motor patterns of ingestion that involve not only release of oromotor patterns, but also modulation of gastric functions.

## Introduction

The performance of ingestive behavior involves actions of both the jaw and stomach. During ingestion of food the jaw is consciously controlled, whereas the stomach is beyond conscious control and instead is subject to modulation by circulating hormones, subconscious reflexes, and chemical signals from the gastric contents. Nevertheless, the functions of the stomach must be coordinated with the jaw during consumption of food. This coordination occurs via reflex and non-reflex responses. Oromotor behaviors like swallowing cause gastric accomodation via a vagovagal reflex^1^. However, coordinated changes in oromotor and gastric function also occur during meal anticipation, suggesting a non-reflex related mechanism to modulate the function of these two systems in concert^2,3^.

We describe here a molecularly identified population of neurons in the central amygdala that both controls bite force and modulates gastric motility and gastric acid secretion. In this way, these neurons can function to coordinate the actions of the jaw with the stomach during the ingestion of food.

## Results

Anatomical tract tracing studies of the central amygdala have found that the medial portion (CeM) of the nucleus is distinct from other sub-regions, namely the CeM makes extensive projections to the brainstem whereas the other sub-regions do not. More recently, spatial transcriptomic analysis of the central amygdala have distinguished the CeM from lateral (CeL) and capsular (CeC) sub-regions by the expression of specific transcription factors^4^. Specifically, the LIM homeodomain transcription factor Isl1 is expressed by neurons in the CeM, but not the CeC or CeL. Given the purported localization of Isl1 expression to the CeM, we expect that central amygdala neurons expressing Isl1 (CeA^Isl1+^ neurons) make substantial projections to brainstem regions.

### Identification of Central Amygdala Neurons Projecting to Ingestive Brainstem Areas

To examine the projections of CeA^Isl1+^ neurons, we injected a virus driving Cre dependent expression of Synaptophysin::mCherry into mice with the Cre recombinase under the transcriptional control of the endogenous ISL1 gene. Viral expression at the injection site matched the anatomical definition of the CeM (Supp. 1A-B). Examination of the synaptic projections from these injections showed that CeA^Isl1+^ neurons made extensive projections to the brainstem and parabrachial areas, as well as a number of other regions (Fig. 1A-D). The brainstem regions receiving projections included parvocellular reticular area (PCRt) a region implicated in control of the jaw, as well as the nucleus of the tractus solitarii (NTS), which is implicated in autonomic control.

**Figure 1:**
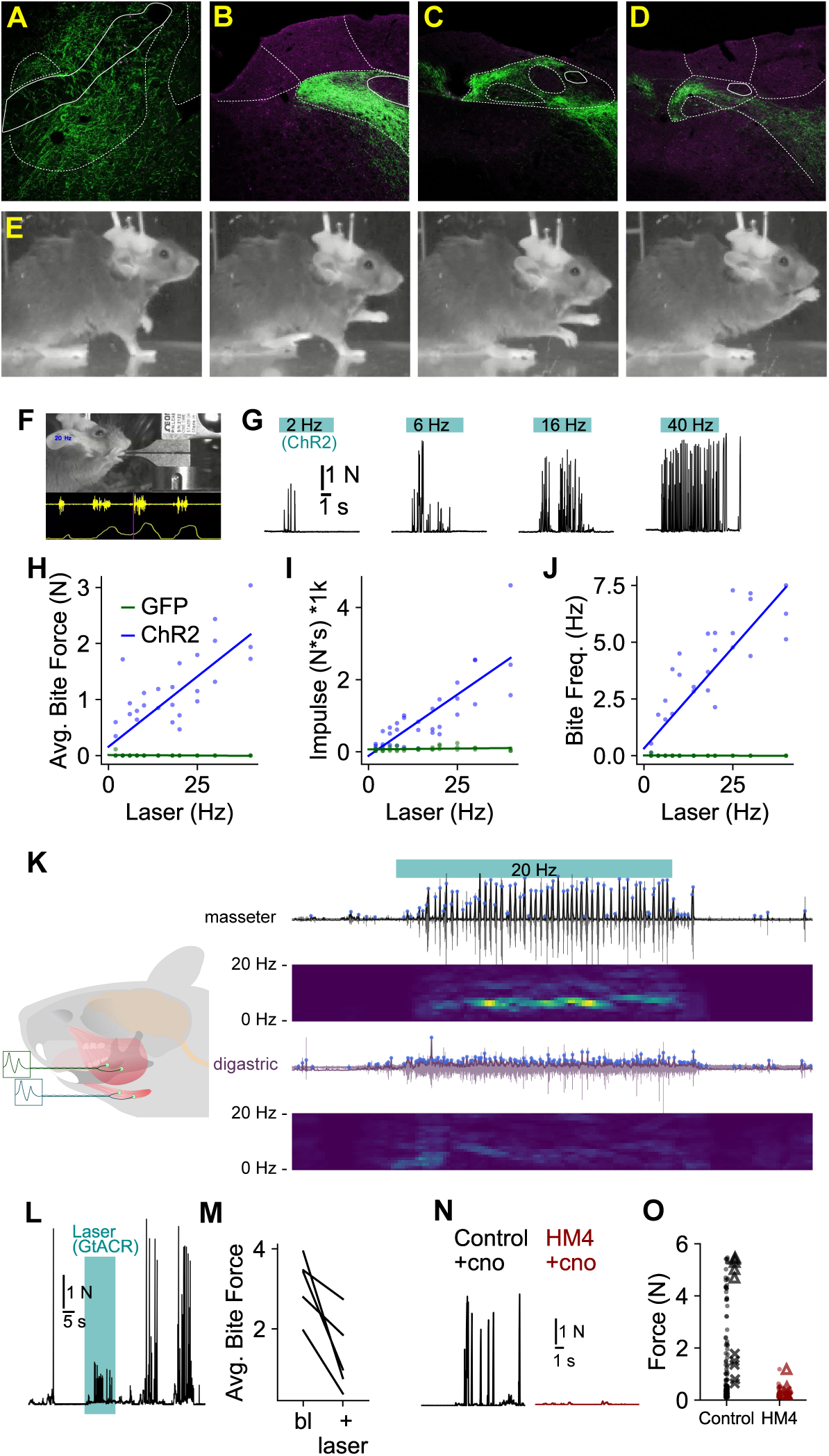
Isl1 neurons are necessary and sufficient to generate high force bites. A-D. Isl1 Neurons project to brainstem regions involved in control of the jaw. Prominent projections are observed to the parabrachial nucleus (PBN, A), the nucleus of the tractus solitary (NTS, B-D), the parvocellular reticular nucleus (PCRt, B-C) and the dorsal part of medullary reticular nucleus (MdD, D). E. Stimulation of Isl1+ neurons in the central amygdala elicited fictive feeding. Sequence of still frames from movie showing grasping of an imagined food object, bringing to mouth and chewing. F. Recording arrangement used to measure stimulation evoked bite force. Animal was placed animal in front of single point load cell, and after they are quietly resting, laser stimulation is commenced. In some cases master EMG activity was also recorded. Here a video record of the session (top image) is synchronized to recorded master EMG (upper yellow trace) and force (lower yellow trace). Vertical magenta line shows time of the frame above, with 0.5 seconds of data preceding (left of line), and 0.5 following (right of line). G. Example traces of load cell force in response to increasing frequencies of CeA^Isl1+^ stimulation. H-J. Stimulation of Isl1+ neurons in the CeA evokes biting behavior. The force (I), number (J) and total impulse (K) of bites evoke by stimulation of CeA^Isl1+^ cells is linearly related to frequency of stimulation (N=3, P=0.00093 bite force, P<0.0001 num. bites, P<0.0001 impulse). Control animals expressing HM4-GFP in CeA^Isl1+^ cells did not bite the load cell with laser stimulation under these conditions. K. EMG recording of rhythmic activation of jaw opening, (digastric) and closing (masseter) muscles jaw evoked by stimulation of CeA^Isl1+^ Neurons. Cartoon on left depicts recording arrangement, traces on right show the rectified integrated EMG. Upper and lower heatmaps show the spectrogram of masseter and digastric activity, respectively. Masseter activity features a prominent peak of activity between 3-8 Hz. L. Chemogenetic inhibition of CeA Isl1 cells reduces bite force. Bite force in an animal expressing ChR2.eYFP in CeA^Isl1+^ neurons (Control), and one expressing the inhibitory DREAD HM4. Both are dosed with CNO. M. Summary data, triangles indicate the maximal bite force and X’s the average bite force, for each animal (N=5). Individual bites are plotted as dots. CNO treatment markedly reduces the mean (t_(8)_=4.70,P=0.0015) and peak (t_(8)_=19.27,P<0.0001) bite force of Isl1-Cre animals expressing HM4 in the CeA. N. Example trace of load cell force before, during and after optogenetic inhibition of Isl1+ CeA neurons. Note that while the laser is on, bite force is reduced. O. Summary bite force data for the optogenetic inhibition experiment (trace in H, N=5), mean bite force is significantly reduced during laser activation (t_(4)_=3.85,P=0.0183).

### Fictive feeding observed on stimulation of CeA^Isl1+^

Based on the pattern of these projections, we hypothesize that the Isl1 neurons may be involved in controlling ingestive behavior. A number of previous reports have described fictive feeding following stimulation of several populations of central amygdala neurons^5^, we tested whether ingestive behaviors or fictive feeding could be evoked by stimulation of these neurons. We injected a Cre dependent virus expressing ChR2, and placed optical fibers over the central amygdala. Activation of CeA^Isl1+^ neurons by laser stimulation evoked feeding related behaviors including behavioral sequences recapitulating the manual or oral acquisition of food and what appeared to be chewing behavior (Fig. 1E). An incidental observation was made during the experiment; in one instance the laser was accidentally activated at a high rate while the experimenter (MHP) was carrying the animal to the recording arena, resulting in the animal biting the experimenter. The bite was substantially harder than expected. This lead us to examine whether the CeA^Isl1+^ may be involved in the generation of high force bites.

### Bite force and masseter EMG are proportional to rate of CeA^Isl1+^ neuron activation

To test whether CeA^Isl1+^ neurons are involved in generation of high force bites, we directly measured bite force during the optogenetic activation of these neurons at different frequencies. To measure force a load cell was modified with a custom pair of bite bar plates^6^. Bite force profiles were recorded with the same system used to drive laser activation and to aquire EMG signals in a subset of animals (Fig. 1F). Isl1^Cre^ mice were injected with AAV driving expression of either ChR2 or GFP (control) in the central amygdala, and optical fibers were placed above the injection site. After recovery, mice were gently positioned in front of the bite plates, and the laser was activated for an 8–10 second period at frequencies ranging form 2–40 Hz. Slice recording of ChR2 expressing CeA^Isl1+^ neurons indicated that neurons could faithfully follow optical stimulation over this range of frequencies (Supp. 1C-D). A single rate of laser stimulation was presented in each trial, and trials were spaced 10–60 seconds apart, during which time the animal was moved away from the bite plates. Peaks in the force trace with prominence greater than 0.3 N were described as separate bites, and all the bites for a single trial were averaged to give an average bite force. In addition, the total trial impulse (area under the force trace) and the bite frequency were averaged for each trial. 2–3 trials at each laser frequency were preformed. Bite measurements were compared between groups expressing GFP and ChR2 with a mixed ANOVA (measure ∼ Opsin X Laser Hz), showing a significant difference for bite force (F_(1,4)_=25.4, P=0.0073), impulse (F_(1,4)_=50.4, P=0.0021), and bite frequency (F_(1,4)_=150.6, P = 0.0002, Fig. 1G-J). There appeared to be a clear correlation between the rate of laser stimulation and the bite force measurements, so we preformed a linear regression analysis between the rate of laser stimulation and the bite force, impulse, and bite frequency. Animals expressing ChR2 showed a significant linear relationship between the rate of laser stimualtion and bite force (r=0.752,P<0.0001), impulse (r=0.813, P<0.0001), and bite frequency r=0.8253,P<0.0001). There were no significant relationships between laser stimulation rate and bite variables in the control group.

We reasoned that this graded relationship between CeA^Isl1+^ stimulation rate and bite force would be apparent at the level of EMG activity in the primary jaw closing muscle, the masseter. Following the measurement of bite force above, the ChR2 group, and one additional animal (N=4), were implanted with EMG wires in the masseter and the digastric, the primary jaw opening muscle. EMG activity was recorded in response to laser stimulation in the home cage (Fig. 1,K). In masseter recordings, the average amplitude of individual EMG bursts, total root mean square amplitude, and the frequency of EMG bursts, all showed a significant positive correlation with the frequency of laser stimulation (Supp. 1E-G). These measures correspond to the force of individual bites, the total trial impulse, and the frequency of bites, respectively. Having established this relationship between stimulation frequency and measurement of masseter EMG activity, we repeated measurements of bite force during CeA^Isl1+^ stimulation, while simultaneously measuring EMG, to examine the relationship between measured bite force and masseter EMG activity. A representative example of this recording is shown in Supp. 1H. Bite force peaks were identified, and the amplitude of masseter EMG activity in a 50 millisecond window preceding the bite peak was measured for each bite (Supp. 1I-K). A linear fit of peak bite force with masseter EMG was highly significant, and very predictive of peak force. These data suggested that masseter EMG could be a reasonable proxy measure for bite force.

### Preferred rhythmic frequency jaw movements is independent of rate of CeA^Isl1+^ neuron activation

A clear aspect of the EMG signal elicited by CeA^Isl1+^ stimulation in the home cage was its rhythmic nature (Fig. 1K). We performed Fourier analysis of the rectified integrated masseter EMG, which showed a prominent peak between 3–7 Hz, the amplitude of which increased with increasing rate of laser stimulation (Supp. 1L-M). Although higher rates of laser stimulation tended to increase the power of the higher frequency portion of the masseter spectrum, they did not appear to shift the frequency of the most prominent peak of spectral activity in the masseter. For laser stimulation rates that reliably evoked rhythmic jaw movements (at or above 6 Hz), we did not observe a significant correlation between the frequency at which the masseter exhibits its greatest spectral power and the rate of laser stimulation (Supp. 1N,U).

Considering the invariance of the fundamental frequency of rhythmic masseteric activity to the rate of laser stimulation, we reasoned that laser stimulation may excite the same central pattern generator active during consumption of food. During biting and chewing of food objects, jaw opening and closing muscles have a stereotypical anti-phase organization to their activity. If laser stimulation of CeA^Isl1+^ drives oromotor movements via the same patterning circuits, we would expect to observe the similar anti-phase relationship between the masseter and digastric. To test this, in a new group of 5 animals, we measured EMG activity of the masseter and digastric during consumption of hard food, soft food, and during laser stimulation of CeA^Isl1+^ neurons. From these EMG records, we compared the spectrum and phase relationship of activity observed during food consumption to those of activity evoked by stimulation of CeA^Isl1+^ neurons. Example EMG traces of the masseter and digastric during laser stimulation, eating chow, and eating Reese’s peanut butter are shown in (Supp. 1O-Q). Masseter EMG activity elicited by laser stimulation had a spectral peak at 3.88±0.31 Hz, not significantly different than the peak observed during consumption of chow 3.50±0.61Hz, or while eating Reese’s peanut cups 3.61±0.36Hz, (ANOVA F_(2,8)_=1.02, P=0.403). In contrast, laser evoked activity of the digastric muscle showed a spectral peak that was significantly different from that observed during consumption of chow or peanut butter. (F_(2,8)_=4.96,P=0.04). Post hoc tests showed that peak frequency in response to laser activation 6.19±0.36Hz is significantly greater than that observed during consumption of peanut butter 4.16±0.48Hz (t_(4)_=2.99,P=0.02, Supp. 1R-T). The phase relationship of masseter and digastric activity was assessed by fitting a damped cosine function to the cross-correlogram of the rectified integrated masseter and digastric EMG signals, representative data and fit, Supp. 1V. Rhythmic EMG activity observed under all conditions exhibited a clear anti-phase relationship between the masseter and digastric (Supp. 1W). Phase relationships during consumption of chow 163.88±6.8 degree and peanut butter 163.53±7.81 degree, or evoked by laser stimulation 169.95±3.48 degree were not significantly different (F_(2,8)_=0.227,P=0.8).

### Inhibition of CeA^Isl1+^ neurons substantially reduces bite force

These data suggested two roles for CeA^Isl1+^ neurons, one is to generate high force bites, and the other to activate the central pattern generator for repetitive jaw movements. We next sought to test whether the activity of this cell population was necessary for these two roles, using both optogenetic and chemogenetic inhibition. Isl1^Cre^ animals were injected with AAV to express the inhibitory opsin GtACR2 in the central amygdala (N=5), allowing us to inhibit CeA^Isl1+^ neurons with laser light. First we measured the ability of animals to generate high force bites when restrained (Fig. 1L-M). Without inhibition, animals generated an average bite force of 3.11±0.34 N which was significantly reduced to 1.34±0.42 N with inhibition of CeA^Isl1+^ neurons (t_(4)_=3.854, P=0.018). Inhibition by GtACR2 did not significantly change the number of bites 19.8±5.6 without inhibition, as compared to 14.8±4.4 during laser activation (t_(4)_,P=0.613). We performed a similar experiment with chemogenetic inhibition (Fig. 1N-O). Isl1^Cre^ animals were injected with AAVs driving either the expression of the inhibitory DREADD hM4Di (N=5) or ChR2.eYFP (N=5). Following injection of CNO, average bite force was significantly lower in the hM4Di group, 1.35±0.42 N as compared to the eYFP group 3.11±0.34 N, (t_(8)_=4.7,P=0.0015). In addition, the number of bites was fewer in the hM4Di group 11.2±3.6 as compared to the control group 35.0±6.75,(t_(8)_=3.11,P=0.014).

### Licking behavior observed with lower rates of CeA^Isl1+^ stimulation

In the course of doing these experiments with the bite meter, we observed that animals would infrequently lick the bite plates on the bite meter, rather than bite them. This raised the possibility that CeA^Isl1+^ neurons may be involved in licking behavior, in addition to biting behavior. To see if this is the case, we removed the bite plates, and substituted a dry sipper (Supp. 1X). The sipper was positioned front of a camera synchronized with the data acquisition system driving the laser stimulation, and recording EMG activity. In contrast to the responses observed following CeA^Isl1+^ in front the bite meter, the presence of the dry sipper lead to reliable observations of licking in response to laser stimulation at rates between 2–6 Hz. There was significant interaction between laser stimulation rate and behavior observed (F_(4,12)_=18.47,P<0.001). At the highest laser stimulation rate, 20 Hz, biting 46±2.74 (count) supplanted licking 0.5±0.5. In contrast at the 4 Hz laser stimulation rate, animals more often licked 12.0±5.31 than bit 2.25±1.93 the sipper (Supp. 1Y).

### A subset of CeA^Isl1+^ neurons’ firing frequency is proportional to bite force in an operant biting task

The preceding experiments suggest that CeA^Isl1+^ cells are involved in generating high force bites. Inhibition experiments suggested that these cells are necessary for high force defensive bites, but do not provide any information as to whether they are active during the generation of high force bites in an appetitive setting. Because the stimulation of CeA^Isl1+^ cells seems to elicit ingestive behaviors, we reasoned that these neurons may be active during biting in an appetitive operant task. To test this prediction, we designed the ‘gobstopper task’ an operant biting task that aimed to mimic normal use of the jaw during feeding. In brief, hungry animals had to bite the force meter in order to receive an intra-oral infusion of a reward solution. Animals were instrumented with chronic intra-oral catheters, and the detection of bites on the force meter exceeding the threshold triggered the solenoid which gated a pressurized reservoir of fluid. A single bite exceeding the threshold lead to a short latency (∼ 100 milliseconds) infusion of ∼10 microliters of reward. Once animals learned to bite the force meter for reward infusion, they were moved to a progressive force threshold, so that after earning 8 rewards, the force threshold would increment by 0.3 Newtons. The bite plates of the force meter were positioned in the operant box closer to the floor, leading mice to make incisal bites. Prior to training, Isl1^Cre^ mice were injected with an AAV driving Cre dependent expression of ChR2, so that CeA^Isl1+^ neurons could be identified by the optotag method. Next a micro-drive bundle of tungsten microwires was implanted above the CeA, and slowly lowered until optically activated units were found. An example of a mouse performing the task is shown in Figure 2A-C. Over the course of the recording session, the amount of force the animal bites with increases to track the force threshold.

**Figure 2:**
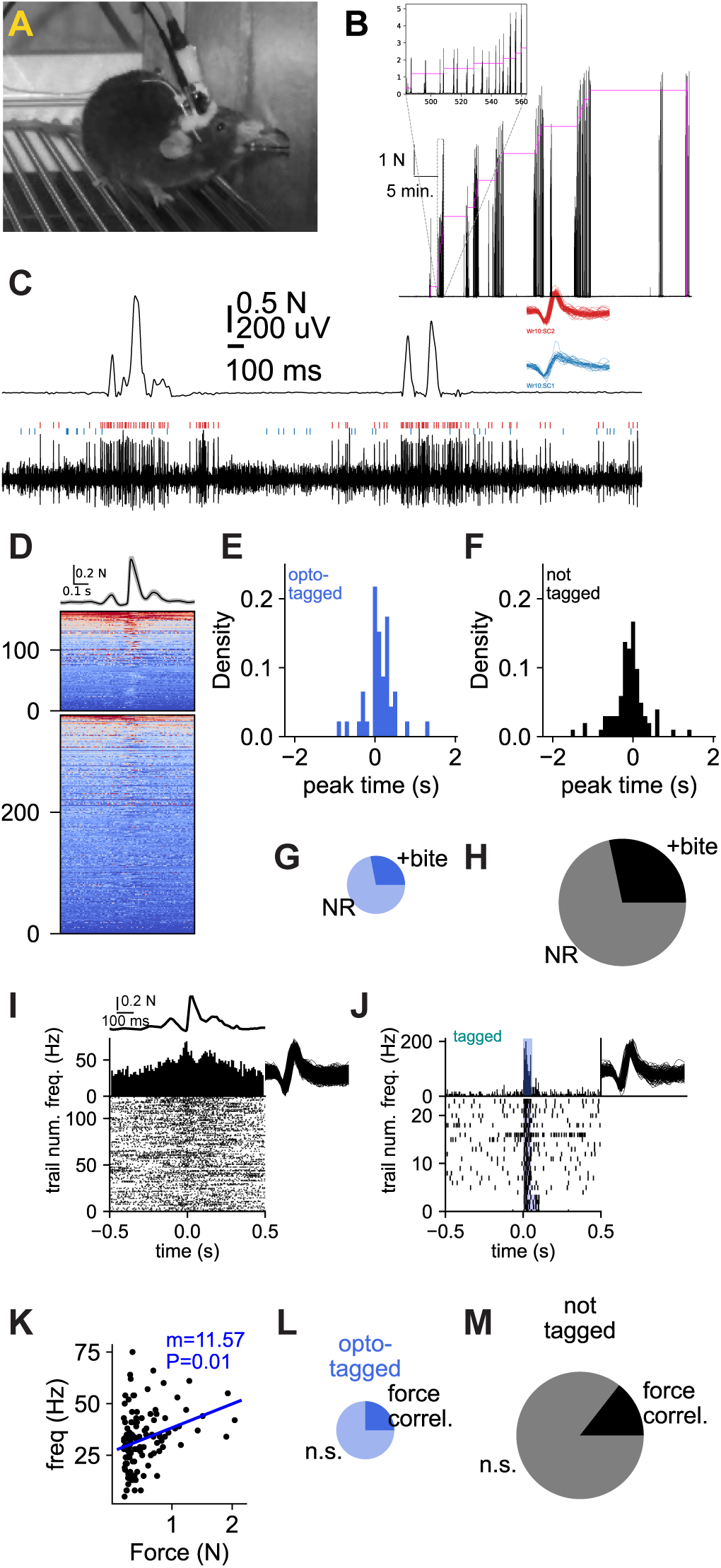
Isl1 cells are active during biting, and their firing rate is correlated with bite force. (A) Representative image showing mouse biting force meter for an intra-oral infusion of lipid reward during opto-tag recording sessions. (B) Example trace of bite force over the course of a recording session with a progressive increase in the reward threshold. The black trace shows the force of individual bites, and the escalating magenta stair-case shows force threshold. (C) Example trace of a unit recording during biting. Upper trace is the force recorded by the bite meter. Lower trace is electrical potential recorded from the tungsten microwire. Inset traces in red and blue are the waveforms of units sorted from this wire. The red unit is active during biting, this unit is in panels F-H. (D) Heat maps showing Z transformed firing rates from 524 cells recorded from 6 mice over 24 sessions. Top trace is the average bite force aligned to the point where it exceeds 0.3 N, the minimal threshold to earn a reward at the beginning of a session. The upper heat map is generated from opto-tagged units (N=164). The lower heat map from non-tagged units (N=360). Impression during recording was that high frequency firing preceded higher amplitude bites. (E-F) Histogram of the times at which the largest changes in firing rates were measured for cells that were significantly active around bites, as measured using the Zeta test. Times of zeta peaks for Opto-tagged cells, are shown in E, non-tagged cells are shown in F. (G-H) Proportion of tagged (G) and non-tagged (H) recorded neurons that showed significant firing rate changes by the Zeta test. (I) Raster plot representative of an opto-tagged CeA^Isl1+^ cell who’s firing rate is correlated with bite force. Time is aligned so that minimal force threshold crossing is at time zero. Upper trace shows average of bite force for all aligned bites, histogram shows the average instantaneous firing rate across all events, raster below has one row for each bite. Rows of raster are ordered from lowest (bottom) to highest bite force. The firing rate of the cell in the between 500 ms before to 20 ms after the threshold crossing is averaged for each bite, and is used for the correlation analysis. (J) Example cell response to laser. (K) Correlation between firing frequency and force of each bite, for raster data in panel I. (L-M) Cells were split into to groups, those showing a significant correlation between firing frequency and the force of each bite, and those that did not. The proportion of opto-tagged Isl1+ is significantly greater among those units who’s firing rate is correlated with bite force, comprising 44% of these force related units, than with units uncorrelated with bite force, comprising 28%, a significant difference (fishers exact 1.91, P=0.011).

A substantial portion of neurons in the CeA appeared to change their activity as the mice bit the force meter. CeA^Isl1+^ neurons had appeared to fire at a faster rate in the period immediately preceding bites to the force meter. To examine this in greater detail, we constructed bite triggered rasters of neurons Z transformed firing rates, Fig. 2D. Manual examination of rasters from individual units gave the impression that changes in neuron firing sometimes exhibited complex dynamics with respect to bite force. To identify units that were significantly modulated during biting, we preformed the Zeta test, as this is agnostic of the timing of changes in neuron firing. Spiking of each unit was aligned to the time that bite force crossed the initial threshold (0.3 N) for all the bites of force meter, and compared to spiking for randomly chosen times. The zeta test identified 46 units out of 164 optotagged cells that exhibited significant firing changes with respect to biting of the force meter. The proportion of bite modulated units among optotagged cells was not significantly different from that observed among the non-tagged cells (102 out of 360, fisher’s exact=0.98, P=1.0, Fig. 2G-H). The zeta test also identifies the time of the most significant change in firing rate, for those significantly modulated units, so we compared the time of the firing rate peak change between optotagged and non-tagged units. The peak time of optotagged units 156±71 ms after threshold crossing, not significantly different from non-tagged units 290±84 ms, (t_(107)_=1.2,P=0.22, Figure 2E-H).

Despite the similar ratio of units that are modulated about the initiation of biting, between non-tagged and optotagged units, we hypothesized that there may be a sub-population of CeA^Isl1+^ neurons that are specifically involved in generating high force bites. Our rationale is that increasing the laser stimulation rate of CeA^Isl1+^ neurons increases the force of bites, so a similar process may occur during spontaneous bites. To examine whether this is true, we measured the peak bite force of each bite, and the firing frequency of each unit in the 200 milliseconds bracketing each peak in bite force. We then tested whether a linear correlation existed between neuron firing rate and bite force for each recorded unit. An example neuron with a positive correlation between firing rate and bite force is shown in (Figure 2I-K). The number of neurons having a significant positive correlation between firing rate and bite force is significantly greater among the optotagged units, (fisher’s=1.97,P=0.004), suggesting these neurons are preferentially involved in generating high force bites (Fig. 2L-M).

### CeA^Isl1+^ neurons active during operant biting are active during food consumption

In addition to recording CeA^Isl1+^ neural activity when biting the force meter, in some sessions, chow was introduced into the operant box at the end of the session, and neural activity was recorded as animals ate (Supp. 2A-H). Neurons that fired at more than 3 standard deviations above their mean frequency during consumption of chow were considered as significantly responding to food. For the sake of comparison, the same significance measure was used in this analysis to determine significant responding during biting of the force meter. By these measures, of the 121 optotagged units that also had recordings of activity during feeding, 16 of these cells were highly active (greater than 3Z) during biting the force meter (Supp. 2H). Among the bite active optotagged cells nearly all (13/16, 81%) were also highly active during consumption of food. In contrast, making the converse comparison, we see not all neurons active during food consumption are also active during biting of the force meter. 50 optotagged units were highly active during consumption of food, and among these 37 (74%) of them were not significantly active during biting of the force meter.

In addition to examining how CeA^Isl1+^ neurons are active during biting, we were curious to see to what extent these neurons maybe involved in licking (Supp. 2I-Q). Because we could evoke licking behavior under some circumstances with lower frequency stimulation and an appropriate visual target, we hypothesized that CeA^Isl1+^ neurons maybe involved in volitional licking behaviors. In a number of ‘gobstopper’ recording sessions performed with a fixed force threshold (1.0 N), we used an open-source contact based potentiometric lick-o-meter to measure licking during electrophysiological recordings. In the same recording session, mice also bit the force meter, so we could compare neural activity during licking and biting with high temporal resolution.

We examined licking related neural activity by two means. In the first we tested whether neurons’ firing rates change significantly in the initiation of a bout of licking (Supp. 2I,L,N). Second we tested whether neurons’ firing is significantly modulated about the generation of individual licks (Supp. 2O,Q). Neuron firing during bouts of licking was compared to firing by the same unit during ‘bouts’ of biting the force meter. Biting bouts were defined as a set of three discreet peaks preceded by a period of 4 seconds with no biting activity. Of the total number of neurons highly active during licking (61) or biting (34). A small number (9) of neurons were active during both bouts of licking and bouts of biting, and they were all optotagged. In contrast, when examining significant modulation of neuron firing frequency during individual licks or bites, almost no overlap was observed. A total of 20 cells were highly active over the course of an individual lick cycle, of these 9 were optotagged. 20 cells highly active over the course of individual bite, of these 19 were optotagged. Only 1 cell was significantly active during both individual licks and bites.

### CeA^Isl1+^ neurons generate bite force by modulating the periodontal­masseteric reflex

In addition to expression of Isl1 in the central amygdala, the transcription factor is also expressed in spinal sensory neurons, and some cranial nerve motor neurons. Unexpectedly, when confirming histologically the viral expression of ChR2 injected in the CeA of an Isl1^Cre^ mouse, eYFP expressing cell bodies were observed in the region of the parabrachial nucleus. This mouse had been left following AAV injection for longer than was planned (6 months). Based on the pattern of spread to areas expressing Isl1 near the injection site (which was limited), we concluded that the spread of this virus must have occurred by trans-synaptic means. Examination of the Allen Brain Atlas showed that the Mesencephalic Trigeminal Nucleus (Me5) cells expressed Isl1 highly, as might be expected due to their similarity to spinal sensory neurons.

To examine whether a synaptic connection existed between CeA^Isl1+^ neurons and Me5 we took advantage of the fortuitous expression of Cre in Me5 neurons to preform rabies virus tracing. To visualize CeA^Isl1+^ cells, we crossed a genetic reporter mouse with the Isl1^Cre^ mouse line. SUN1::Isl1^Cre^ mice were breed, that expressed GFP in CeA^Isl1+^ neurons. In these mice, a Cre dependent AAV expressing rabies G protein was injected into the Me5 nucleus. Three weeks after this, ΔG rabies virus expressing RFP was injected at the same coordinates in Me5, and in the same surgery, cholera toxin b conjugated to Alexa647 was injected into either the NTS or the PSTn (Fig. 3A). Seven days after injecting the rabies, animals were perfused for histology. Starter cells were tightly localized in Me5 (Supp. 3A). Cells in the CeA were categorized by RFP expression (projecting to Me5 neurons), SUN1 expression (CeA^Isl1+^ neurons), and Alexa647 signal (projecting to NTS or PSTn according to the surgical case). All RFP+ neurons in the CeA projecting to Me5 expressed the SunTag. We conclude that CeA^Isl1+^ neurons specifically project to Me5. For the two cases where ctb was injected in the NTS, only a single CeA^Isl1+^ neuron projecting to Me5 was positive for ctb. This suggesting that very few CeA^Isl1+^ neurons make divergent or collateral projections to both the NTS and to Me5. A similar result was observed for the two cases where ctb was injected into the PSTn (No ctb+ and RFP+ cells). We conclude that there are subgroups of CeA^Isl1+^ neurons projecting to different brainstem regions, and almost no CeA^Isl1+^ cells with divergent projections to multiple regions.

**Figure 3:**
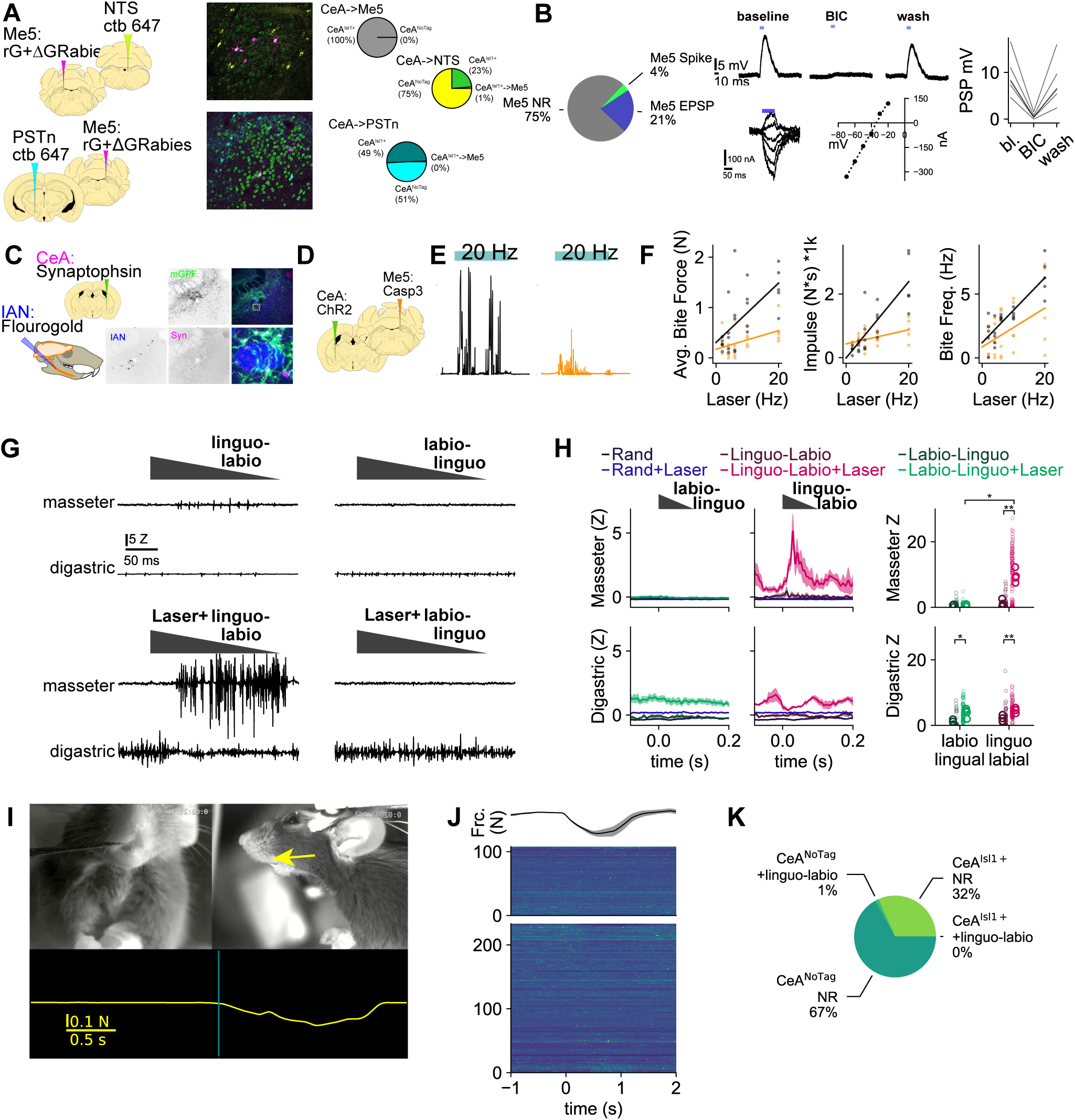
CeAIsl1^+^ neurons increase bite force by amplifying the periodontal masseteric reflex. A. Rabies virus tracing was preformed to check the putative connection from Isl1+ neurons to Me5 neurons. Fortuitously, Me5 neurons express Isl1, allowing the experiment to be preformed using Isl1^Cre^ mice. To mark the Isl1 neurons in the central amygdala, the nuclear INTACT reporter mouse was crossed with the Isl1 Cre line. AAV1 split TVA-Rabies G was injected into the caudal Me5 of INTACT::Isl1^Cre^ mice. Two weeks later, ΔG rabies virus expressing mCherry was injected into Me5. In the same surgery ctb 647 was injected into either the NTS (N=2), or the PSTn (N=2). All rabies positive cells identified in the central amygdala where positive for the INTACT nuclear GFP staining. A single NTS projecting CeA neuron was positive for rabies. B. Patch recording of Me5 neurons. In response to activation of CeA^Isl1+^ terminals we observed EPSPs or action potentials in Me5 neurons (pie plots). Upper traces show depolarizing PSPs recorded in an Me5 neuron. LED activation of ChR2 terminals indicated by blue box above each trace. Left upper trace, depolarizing PSP in normal aCSF. Middle, addition of bicuculline reduces PSPS by 94%. Right trace, washout of bicuculline recovers PSP, at 82% of its initial amplitude. Lower family of traces shows post-synaptic currents measured at holding potentials between −70 and −20 mV. The reversal potential of the PSP was calculated in 6 cells, and averaging −34.1 ± 5.4 mV. C. The synaptic projections of Isl1+ CeA neurons to tooth sensory neurons were examined by injecting a AAV1-mGFP:Synaptophin_mRuby into the CeA of Isl1Cre animals, and the retrograde tracer fluorogold was injected into the inferior alveolar nerve. GFP and mRuby were clustered around the cell bodies of FG labeled Me5 neurons, suggesting a synaptic connection between CeA^Isl1+^ neurons and tooth sensory neurons. D. Cartoon shows injection scheme: AAV5 DIO ChR2 was injected into the CeA, and fibers were placed above the CeA, and in the same surgery, AAV1 Casp or AAV1 HM4 (control) was injected into Me5. Three weeks following bite force was measured for each group. E. Representative bite force traces recorded when animal bites the load cell in response to 20 Hz stimulation of CeA Isl1 cells in control (black) and caspase lesioned (orange). F. Comparison of laser evoked biting of the load cells between the caspase and control animals shows a reduction in the average force of each bite (left panel, F_(1,8)_=6.34 P = 0.036) further a significant interaction between Group x Laser frequency was observed (interaction F_(4,32)_=3.53, P=0.017). Post hoc tests confirmed that bite forces observed following 2 Hz stimulation did not differ significantly (0.35 ± 0.10 N Control v. 0.29 ± 0.04 Caspase, t_(8)_=0.55,P 0.592), whereas bite force observed during 20 Hz stimulation was significantly less in the caspase group (0.56 ± 0.125 N) as compared to control (1.45 ± 0.23 N, t_(8)_=-3.70, P=0.0061). A similar pattern of significance was observed for the impulse (total integrated force), (middle panel, interaction, Group X Hz, F_(4,32)_=8.21, P = 0.0001), No significant difference was observed in the number of bites between the control and caspase groups (right panel, F_(1,8)_=1.77,P=0.219). G. Representative traces of jaw muscle activity evoked by stimulation of the incisor teeth, a measurement of the periodontal jaw closing reflex. Traces on the upper row are muscle activity evoked by tooth stimulation alone, without laser stimulation. Lower traces show muscle activity evoked during simultaneous tooth stimulation and laser activation of CeA^Isl1+^ cells. H. Summary data of muscle activation evoked by tooth stimulation and laser activation. Line plots are integrated rectified muscle amplitude aligned to stimulation time and averaged. Stimulation in the linguo-labio direction evokes a small activation in the masseter, which is dramatically potentiated by Isl1 activation. Dot plots are the maximal signal for each trial (small dots) and the average of all trails for each mouse (larger dots) N=4 mice. Upper plot show amplitude of the masseter response, and lower plot shows the amplitude of the digastric response. A mixed ANOVA, comparing EMG amplitude as a function of the interaction between muscle identity, laser state, and stimulus type show significant interactions between the three variables (F_(3,9)_=104.4, P<0.0001). Post-hoc tests showed that laser activation significantly increases the amplitude of the masseter contraction evoked by stimulation of the teeth in the linguo-labio direction, from 1.07 ± 0.23 to 9.45 ± 0.75 (AU), P=0.001, but has no effect on amplitude of master contraction evoked by stimulating the teeth in the labiolinguo direction 0.61 ± 0.11 laser off vs 0.52 ± 0.077 laser on, P=0.586. In contrast, laser activation significantly increased the amplitude of digastric contraction evoked by stimulation of teeth in either the lingo-labio or labio-linguo direction. I. Video visualization of experimental set up to measure single unit activity in the central amygdala during stimulation of the teeth. Example of stimulation of the incisor teeth in the linguo-labio direction. Upper frames show synchronized head-on and side view video recording. The yellow arrow on the right video frame indicates the direction of force applied to the incisors. The probe was pressed against the lingual surface of the teeth. Lower yellow trace shows the force applied by manual stimulation to the incisor, downward deflection indicates the linguo-labio direction. The vertical cyan bar transecting the force trace indicates the time of the above video frames. J. Summary heat map data showing single unit responses to stimulation of incisor teeth in the linguo-labio direction. Black trace above the heatmaps shows the average force applied across each recording site, gray shading indicates standard error of the mean. 341 cells in total were recorded from three mice. Four site were recorded in each mouse, for a total of 12 distinct recording locations. Of the recorded cells, 108 were classified as Opto-tagged by the SALT criteria, they are labeled as CeA^Isl1+^. The remaining 233 cells are labeled as CeA::NoTag. The upper heat map shows responses of the CeA^Isl1+^ cells to the tooth stimulation, the lower heat map shows the CeA::NoTag cells. Colors in heat map span from +3Z (yellow) to −1.5 Z (dark blue). K. Pie plots showing proportion of CeA^Isl1+^ and CeA::NoTag that significantly respond to stimulation of incisors in the linguo-labio direction.

To further test whether there is a synaptic connection between CeA^Isl1+^ neurons and Me5 neurons, we performed acute brain slice recordings from Isl1^Cre^ mice injected with a Cre dependent AAV to drive ChR2 expression in CeA^Isl1+^ neurons. Then brain slices were taken from the parabrachial region. Me5 neurons were identified by their size, location, and intrinsic electrical properties, namely: a prominent I_h_ current, a low input resistance and voltage dependent membrane resonance around 100 Hz (Supp. 3B). Recording from 80 Me5 neurons in 5 mice, we found 20 Me5 neurons that were excited by CeA^Isl1+^ terminal stimulation. Of the 20 excited neurons, 17 showed depolarizing EPSPs, and 3 had spiking responses to CeA^Isl1+^ terminal stimulation (Fig. 3B). In a subset of these recordings, we washed bicuculline into the bath, which reduced optically evoked EPSPs from 12.45±0.92mV to 0.83±0.15mV. Extended washout partially recovered the optically evoked EPSP to 10.0±1.04mV. For the three Me5 neurons showing spiking responses to CeA^Isl1+^ stimulation, we were unable to complete the bicuculline wash in and wash out. In addition in another set of 6 Me5 neurons showing optically evoked EPSPs, we measured the reversal potential of the PSP by measuring post-synaptic currents at holding potentials between −70 and −20 mV. The reversal potential of the PSP −34.±5.4 mV.

The relatively low proportion of responsive Me5 neurons to optical activation of CeA^Isl1+^ terminals suggested to us that there may be a selective type of Me5 neuron receiving input from the CeA^Isl1+^ cells. Neurons in the Me5 can be segregated into two functional and anatomically distinct groups. The majority are muscle spindle sensory neurons, projecting to muscle spindles in the masseter muscle, and sensing changes in its length. A smaller portion, roughly 20%, project to the periodontal ligament, specifically the lingual portion of the ligament around the incisor teeth. We hypothesized that CeA^Isl1+^ cells may be making synaptic contacts with Me5 neurons projecting to the periodontal ligament. To test this we injected into CeA of Isl1^Cre^ mice a Cre dependent AAV expressing eGFP and the fusion protein synaptophysin::mCherry, to mark presynaptic terminals. Three weeks following this, we then injected FluoroGold into the Inferior Alveolar Nerve, to retrogradely label Me5 neurons that project to the periodontal ligament. A number of mCherry puncta, putative synaptic contacts, were observed over the cell bodies of Me5 neurons (Fig. 3C).

Because Me5 neurons make monosynaptic excitatory connections with jaw closing motor neurons, and CeA^Isl1+^ neuron appear to excite them, we reasoned that these Me5 cells may be important in the generation of forceful biting observed with CeA^Isl1+^ stimulation. To test this prediction, we again took advantage of the fact the Me5 neurons express Isl1, and using Isl1^Cre^ animals, we injected Cre dependent AAVs in the CeA and Me5 regions to drive cell type specific expression in either area. We lesioned the Me5 neurons by injecting Cre dependent caspase into the Me5 region of Isl1^Cre^ mice. In the same surgery, Cre dependent ChR2 was injected into the CeA, and optical fibers were placed above the CeA region, to excite CeA^Isl1+^ neurons, (Fig 3D-F). Control animals were injected with hM4Di, but did not receive CNO. Comparison of laser evoked biting of the load cell between the caspase and control animals shows a reduction in the average force of each bite (F_(1,8)_=6.34, P=0.036 Fig. 3E-F). Further, a significant interaction between Group x Laser Frequency was observed (interaction F_(4,32)_=3.53, P=0.017). Post hoc tests showed that bite forces observed during 20 Hz stimulation were significantly less in the Caspase group (0.56±0.125 N) as compared to control (1.45±0.23 N, t_(8)_=-3.70, P=0.0061). In contrast, 2 Hz stimulation average bite force did not differ significantly between control 0.35±0.10N and caspase groups 0.29±0.04N,t_(8)_=0.55,P=0.592. A similar pattern of significance was observed for the impulse (total integrated force) recorded during the period of laser stimulation. (interaction, Group X Hz, F_(4,32)_=8.21, P=0.0001). Post-hoc pairwise tests again showed that at 20 Hz stimulation, impulse measured in caspase group 893.86±158.58 was significantly less than in the control 2546.48±376.86 N*s, t_(8)_=-3.49,P=0.008. In contrast, at 2 Hz stimulation, the Caspase group actually had slightly greater impulse than control (474.37±13.53 vs. 272.97±48.58 N*s, t_(8)_=4.04,P=0.0037). No significant difference was observed in the number of bites between the Control and Caspase group (F_(1,8)_=1.77,P=0.219).

This finding suggested that CeA^Isl1+^ cell activity may be modulating Me5 activity to generate jaw closing force. One possible mechanism by which this could occur is that CeA^Isl1+^ activation could increase the gain of a jaw-closing reflex. The periodontal-masseteric reflex is a jaw-closing reflex that has been described in cats, dogs, and rats. This reflex is directionally specific, presumably due to the specific projection of Me5 sensory into the portion of the periodontal ligament ab-lingual to the incisors in rats, or canines in cats and dogs. Deflecting the incisor teeth in the linguo-labio direction will excite the periodontal Me5 neurons, whereas deflections in the labio-linguo direction does not.

To test whether CeA^Isl1+^ neuron activation modulates the periodontal masseteric reflex, we measured this reflex in urethane anesthetized Isl1^Cre^ mice. Mice (N=4) were injected with Cre dependent AAV driving ChR2 in the CeA. Four weeks after viral injection, animals were instrumented with EMG wires in the masseter and digastric muscle, and a headplate was affixed to the headcap which already held the optical cannula. Animals were anesthetized with Urethane and headfixed in front of a two camera system which synchronized camera frames with EMG records. The incisor teeth were manually probed with a blunted forceps in either the linguo-labio or labio-linguo direction and the resulting activation of the masseter and digastric jaw muscles was recorded. On alternate trials, CeA^Isl1+^ neurons were activated at the same time as the tooth stimulation was preformed. Example traces from a single stimulus presentations are shown in Fig. 3G. A mixed ANOVA, comparing EMG amplitude as a function of the interaction between muscle, laser state, and stimulus type show significant interactions between the three variables (F_(3,9)_=104.4, P<0.0001). Post-hoc tests showed that laser activation significantly increases the amplitude of the masseter contraction evoked by stimulation of the teeth in the linguo-labio direction, from 1.07±0.23 to 9.45±0.75(AU), P=0.001. Laser stimulation had no effect on amplitude of master contraction evoked by stimulating the teeth in the labio-linguo direction 0.61±0.11 laser off vs 0.52±0.077 laser on, P=0.586 (Fig. 3H).

The activation of CeA^Isl1+^ neurons had an effect to dramatically increase masseter muscle activity in response to stimulation of the teeth in a manner that would occur during the initiation of a bite of food. This raised the possibility of a positive feedback mechanism if CeA^Isl1+^ neurons are themselves activated by stimulation of the teeth. We tested whether stimulation of the teeth activated CeA^Isl1+^ cells by performing optotag recordings in urethane anesthetize animals (N=3) while stimulating the incisor teeth (Fig. 3I-K). In these experiments manual stimulation of the teeth was performed in a similar manner as in the periodontal-masseteric reflex experiment, except now the forceps probe was carried on a force meter. This allowed us to examine relationships between force applied to the tooth, and cell activity. To our surprise, out of the 108 optotagged putative CeA^Isl1+^ cells recorded from, none were significantly modulated by stimulation of the teeth in the linguo-labial direction (Fig. 3I-K). Stimulation of the teeth in the labio-linguo direction, or opening of the jaw were equally ineffective at modulating firing of CeA^Isl1+^ neurons (Supp. 3K-P).

Considering that CeA^Isl1+^ stimulation modulated a basic reflex arc, we wondered if an additional means by which CeA^Isl1+^ activation could increase bite force is by suppressing reflexes that would functionally oppose jaw closing, by suppressing jaw opening reflexes. We tested this in the same experiment in which the tooth stimulation was performed. The blunt probe was used to push into the hard palate evoking a jaw opening reflex. This stimulus was presented either with the laser off and on, and the magnitude of digastric EMG activity was compared (Supp. 3C-D). No significant difference was observed in mean digastric activity between laser on and off conditions t_(2)_=-0.002,P=1.0.

Because the jaw opening reflex is a defensive reflex, serving to protect the jaw and oral cavity from injury, we reasoned that CeA^Isl1+^ stimulation may only potentiate ingestive oromotor reflexes. If this is true, we predict that CeA^Isl1+^ stimulation may potentiate other ingestive reflexes. One ingestive reflex which has been described were gentle probing of upper lip area can evoke licking behaviors. Experiments with anesthetized rats and cats have described ingestive responses to lip stimulation particularly when performed concomitantly with electrical activation of the lateral hypothalamus. We tested whether in a similar paradigm stimulation of CeA^Isl1+^ neurons could facilitate licking in response to lip stimulation. In urethane anesthetized mice, gentle touch to the portion of the upper lips close to the mouth could occasionally evoke small protrusions of the tongue. When this stimuli was given while activating the laser to stimulate CeA^Isl1+^ neurons, tongue protrusions that appeared similar to licking were likelier to be observed and larger in amplitude (Supp. 3E-J). The likelihood of evoking a lick on a single stimulation trial increased from 0.11±0.04 to 0.43±0.07 (t_(3)_=-4.69,P=0.0183). In addition, the distance the tongue was protruded increased from 2.43±0.83to 14.8±1.94pixels, (t_(3)_=-4.82,P=0.017). Interestingly, monosynaptic tracing of the inputs to CeA^Isl1+^ cells identified two areas associated with fluid intake, the PVH and SON. Immunofluorescence staining of these sections showed that within these regions, vasopressin cells are projecting to CeA^Isl1+^ neurons, whereas oxytocin cells do not (Supp. 3Q-V).

### CeA^Isl1+^ neuron activity is important for the consumption of hard foods

The activity of CeA^Isl1+^ neurons seems to play an important role in generating forceful closure of the jaw, under defensive and appetitive contexts. Based on the preceding experiments, we expect that these neurons are important for the consumption of hard food that requires high force biting. To test this directly, we observed the effect of stimulating or inhibiting CeA^Isl1+^ neurons while animals consumed a hard food: dried pasta. Isl1^Cre^ animals were injected with AAVs to express either the inhibitory opsin GtACR2 or the excitatory opsin ChR2, and optical fibers were placed over the CeA. Once recovered, pasta eating was measured in food deprived animals (Fig. 4A-D). The latency to finish a 2 cm length of dry pasta is decreased by activation of Isl1 cell activation (ChR2 group) and increased by Isl1 cell inhibition (GtACR) F_(1,8)_=13.05, P=0.00685 [Opsin*Laser]. During the ‘No Laser’ phase of the experiment, the latency to finish a single 2cm length of pasta is not significantly different between animals expressing ChR2 and GtACR2, t_(8)_=0.687, P=0.512. In contrast, during the laser on phase, latency to finish is significantly different between the activation and inhibition group t_(8)_=-3.52, P,=0.0078, having increased in the inhibition group (as compared to the no laser baseline) t_(4)_=-2.78, P=0.0497, and trended to decrease in the activation group t_(4)_=2.42,P=0.073.

**Figure 4:**
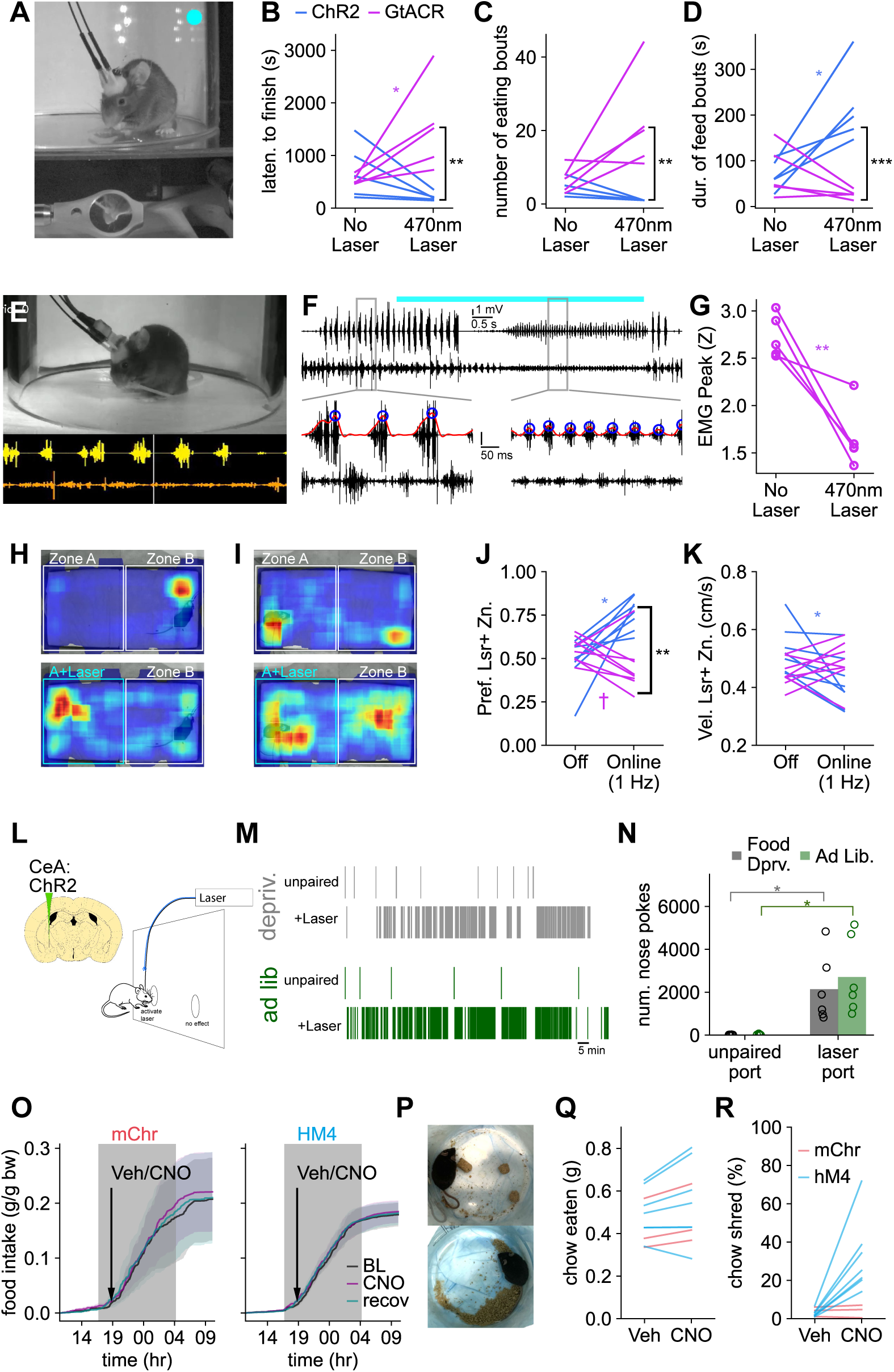
Functional involvement of CeA^Isl1+^ in the consumption of hard foods and motivated behaviors. A-D. CeA^Isl1+^ neurons bi-directionally modulate consumption of hard food, dry-pasta. A. Example frame of video recording during pasta eating, a mirror was placed under the animal to facilitate observation of pasta consumption. B. The latency to finish a 2 cm length of dry pasta is decreased by activation of CeA^Isl1+^ neurons (ChR2 group) and increased by inhibition (GtACR) C. ChR2 stimulation consolidated pasta eating so that it occurred in a single uninterrupted bout. GtACR inhibition had the opposite effect, the consumption of a single piece of pasta was frequently interrupted, such that more feeding bouts were required to finish. D. The average duration of each feeding bout was increased by Isl1 cell activation, and reduced by inhibition. E. Following these feeding measurements, animals in the GtACR group were implanted with EMG wires in the masseter and digastric to examine the effect of CeA^Isl1+^ cell inhibition. Frame shows synchronized EMG and behavior recording, traces below still show masseter (yellow), and digastric (orange). F. Example traces of masseter and digastric EMGs, when CeA^Isl1+^ neurons are inhibited starting after initiation of pasta eating. Top trace, masseter, lower trace digastric, pull out details show the masseter burst detection (blue circles), and amplitude measurement (red enveloping line). Cyan bar above top trace indicates period during which laser light is constantly active. G. Summary data, CeA^Isl1+^ inhibition reduces the amplitude of masseter EMG bursts during feeding t_(4)_=5.27, P=0.0062 H-K. Activation or inhibition of CeA^Isl1+^ neurons alters the location preference of animals in a real-time online place preference assay. H. Heatmaps showing the location preference for an animal expressing ChR2 in CeA^Isl1+^ neurons, before the activation of the laser rule (upper heat map) and again after the laser is conditionally activated (lower heat map) when the animals is present in Zone A. The laser is pulsed at a relatively low frequency, 1 Hz, that is not seen to evoke oromotor behaviors. I. Similar to H, but in an animal expressing the inhibitory opsin GtACR in CeA^Isl1+^ neurons. In the lower heat-map, the laser is continuously on while the animal is in Zone A, as compared being pulsed at 1 Hz while the animal is in Zone A in Panel H. For both of the lower heat maps in H and I, the laser is off while the animal is in Zone J. Summary place preference data for groups of animals expressing GtACR or ChR2. A significant interaction between activation of the laser and opsin type is observed by mixed ANOVA analysis. F_(1,14)_=23.67,P=0.0002. With the laser-place rule ‘online’, a significant difference between preferred side is apparent between the ChR2 and GtACR groups. K. Locomotor speed was measured during the two phases of the experiment. A significant decrease in locomotor walking speed was observed in the ChR2 group.Despite this effect the average velocity between the ChR2 and GtACR groups was not different while the laser rule was online. L-N. Contingent activation of CeA^Isl1+^ neurons acts as an operant reinforcer in a nose-poke self-stimulation paradigm. L. Diagram showing paradigm of self-reinforcement design. Nose-pokes into the laser paired port activated a brief train of stimulatory laser light. Nose-pokes into the other port had no effect. M. Example event plot of poke events for one experimental subject. Experiment was conducted on consecutive days, with animals food deprived on the first day, and then the next day following reintroduction of ad lib food. N. Animals nose-poked into the laser paired port significantly more than the unpaired port. No significant difference was observed between number of pokes into the laser port when animals were food deprived as compared to ad. lib. fed. O-R. Inhibition of CeA^Isl1+^ neurons does not reduce food intake, but effects feeding proficiency. O. Isl1^Cre^ mice expressing either hM4Di (inhibition, N=7) or mCherry (control, N=3) were allowed free access food available from FED3 feeders. Once a stable circadian pattern of intake was established, animals were dosed with Veh-CNO(4mg/kg)-Veh on three successive days two hours after lights out when mice began to consume most of their food. No significant differences in food intake was observed. P-R. Consumption of regular chow food intake was measured in the same animals, in a bedding free arena. Q. Weight of chow consumed was not significantly reduced by CNO treatment, but more of the food was shredded (R).

Stimulating or inhibiting CeA^Isl1+^ cells also bidirectionally changed the feeding bout structure of pasta eating. Inhibition of the cells caused mice to interrupt their consumption more frequently, animals much more frequently dropped the food and stopped eating. Stimulation of CeA^Isl1+^ cells consolidated consumption of every 2 cm length of pasta into a single bout, invariably, mice did not drop or pause their consumption during stimulation F_(1,8)_=8.571, P = 0.0191. Pairwise tests showed significantly more bouts in the GtACR group as compared to the ChR2 group, but only during the laser on phase of the experiment, t_(8)_=-3.54,P=0.0076. Accordingly, the length of individual feeding bouts was increased by stimulation, and reduced by inhibition.

We predicted that if inhibition of CeA^Isl1+^ was reducing bite force during pasta eating, we should observe a reduction Masseter EMG activity, considering the clear linear relationship between EMG amplitude and bite force detailed in Supp. 1I. To test this, after measuring changes in pasta eating, Isl1^Cre^ mice expressing GtACR were implanted with EMG wires in the masseter and digastric (Fig. 4E-G) Recordings of masseter EMG during pasta eating, showed a reduction in amplitude upon inhibition of CeA^Isl1+^ neurons, t_(4)_=5.27, P=0.0062. While inhibiting CeA^Isl1+^ cells, animals were frequently seen to stop eating, dropping the pasta, but continuing to make oromotor movements that appeared subtly distinct from chewing. Inspecting the change in masseter EMG observed during CeA^Isl1+^ inhibition, a qualitatively different pattern of masseter contraction is observed, that has been described as incisal sharpening.

Consumption of hard food is a motivated behavior, accordingly, if manipulating CeA^Isl1+^ activity has an effect to manipulate motivation, then changes in pasta eating observed with these neural manipulations could be due to motivational changes rather than changes in motor capacity. To see if CeA^Isl1+^ neuron activity can modulate motivated behaviors, we tested whether opto-genetically manipulating their activity can introduce a place preference. Manipulating the activity of CeA^Isl1+^ neurons bidirectionally alters the place preference of animals in an optogenetic real-time online place preference assay, (Fig. 4H-K). Animals’ side preference showed a significant interaction between Laser Rule and Opsin type, F_(1,14)_=23.67,P=0.0002. Pairwise tests show that when the laser rule is offline, side preference is not significantly different between the ChR2 and GtACR groups t_(14)_=-1.316,P=0.209. In contrast, with the laser-place rule ‘online’, a significant difference between preferred side is apparent between the ChR2 and GtACR groups, t_(14)_=5.149,P=0.0001. Post-hoc tests show that the preference for the laser paired zone increases in animals expressing ChR2 when the laser rule is online. t_(7)_=-4.57,P=0.0026 and trends toward a decrease in animals expressing GtACR2, t_(7)_=2.07829,P=0.076.

Assessment of behavior in an open field arena suggested that neither stimulation nor inhibition of CeA^Isl1+^ neurons changed the expression of anxiety behavior (Supp. 4G-K). Mice did not shift the duration spent in the center of the open field area with either stimulation or inhibition. F_(1,8)_=1.63,P=0.237. In both open field and place preference arenas, stimulation of CeA^Isl1+^ neurons significantly reduced average velocity. In the place-preference experiment, walking velocity in ChR2 animals between the laser-off and laser-rule on periods slowed from 0.52±0.03 to 0.42±0.3 cm/s, t_(7)_=3.19,P=0.015, and in the open field area the reduction was more apparent slowing from 6.59±0.40 to 2.82±0.26 cm/s t_(4)_=14.73,P=0.00012.

The ability to change place preference by manipulating CeA^Isl1+^ activity suggested that this cell population may influence motivation. To examine this more directly, we tested whether stimulation of CeA^Isl1+^ cells could function as a reinforcer in an operant nose-poke paradigm. A second group Isl1^Cre^ animals (N=6) was used for this experiments, as in previous experiments Cre dependent AAVs expressing ChR2 were injected in the CeA, and optical fibers were placed above the region. Nose pokes to one port were paired with activation of laser to stimulate CeA^Isl1+^ cells, pokes to the other port did not activate the laser (Fig. 4L). Because CeA^Isl1+^ stimulation can evoke fictive feeding, we reasoned that the ‘reward salience’ of this neural activation could be dependent on the animal’s hunger state. Specifically, we predicted that the efficacy of CeA^Isl1+^ neuron stimulation as a reinforcer would be greater in animals that were food deprived.

Laser stimulation of CeA^Isl1+^ neurons was a very effective operant reinforcer, leading to a strong preference to poke the laser paired poke port F_(1,5)_=13.14,P=0.015,(Fig. 4L-N). Further, this reinforcing effect was independent of hunger state, we did not see a significant interaction between port preference and hunger state F_(1,5)_=2.78,P=0.16, interaction: port X internal state. Food-restricted (hungry) mice poked the laser paired port 3291.3±851.7 times in an hour, significantly more than pokes to the unpaired port 19±3.6. Pokes to the laser paired port in refed (sated) mice 3910.7±1028.9 were not significantly different from the number to the paired port in hungry mice t_(5)_=1.71,P=0.15. Lesioning Me5^Isl1+^ neurons in Isl1^Cre^ animals did not seem to suppress the effectiveness of CeA^Isl1+^ neurons stimulation to function as an operant reinforcer (Supp. 4L-O). This suggested to us that CeA^Isl1+^ cells are not lesioned or disrupted by the Cre dependent caspase virus, when injected in the mesencephalic area.

These results suggest the possibility that modulating the activity of CeA^Isl1+^ neurons changes the animals’ motivational drive, and independent of capacity to generate force, this change in motivation alters consumption of pasta. If this is the case, then manipulations that change CeA^Isl1+^ activity should alter food intake, even when the food provided is relatively softer and easier to consume. To test this, we used a chemogenetic strategy to inhibit CeA^Isl1+^ neurons and measured food intake following CNO injection. Animals were feed small pellets softer than the standard lab chow (Fig. 4O). The smaller pellets can be chewed without incisal biting. Pellets were dispensed from an automated feeder (FED3), and mice were housed without cage mates to allow accurate measurement of food intake. Isl1^Cre^ animals expressed either ChR2.mCherry (N=3, control), or hM4Di.mCherry (n=7). Food intake was binned into 10 minute intervals and normalized by body weight. Two hours after lights off, at which point animals begin to consume food, animals were injected either with vehicle (saline) or CNO 4mg/kg. No significant difference in food intake is observed with CNO treatment between groups, F_(1,8)_=0.023, mixed ANOVA on change from baseline, P=0.883. Because we do not see a change in food intake with inhibition of CeA^Isl1+^ neurons, this suggests that these neurons are not a critical for maintaining motivation to consume food in hungry animals.

While acute CeA^Isl1+^ neuron inhibition interferes with consumption of pasta, it does not appear to suppress motivation, or capacity to eat small pellets of foods. Because pasta eating is a specific skill that mice learn and improve their performance on, it could be that the interference with pasta eating observed with optogenetic CeA^Isl1+^ inhibition is related to compromising the performance of a specialized learned motor skill, rather than diminishing the capacity to generate forceful bites. If this is the case, then presumably the performance of ingestive motor behaviors that are learned since weaning should be less sensitive. Meaning that inhibition of CeA^Isl1+^ neurons should not interfere with consumption of the standard laboratory chow. To test this, we used the same animals to test how their ability to eat standard chow is effected by inhibition of CeA^Isl1+^ neurons.

Inhibition of CeA^Isl1+^ neurons caused animals to shred their food much more. On observing this tendency, we moved animals to a hard-bottom arena, so that we could collect all the uneaten food. We quantified food shredding by measuring the percentage of uneaten food that could pass through a coarse sieve (Fig. 4P-Q). Inhibition of CeA^Isl1+^ neurons significantly changed food shredding, F_(1,8)_=5.77,P=0.043, interaction AAV X i.p. injection. Post-hoc tests confirmed that the hM4Di group shredded 32.16±7.3% of their food, significantly more than the control 4.0±1.93, t_(8)_=2.41,P=0.042, after CNO treatment. Vehicle injections did not significantly change shredding t_(8)_=0.67,P=0.52. CNO injection did not affect the amount of food consumed F_(1,8)_=0.01,P=0.9, control and hM4Di animals ate 0.55±0.07 and 0.47±0.08 grams of food respectively.

### CeA^Isl1+^ neuron activity modulates gastric acid secretion

Because the consumption of food necessarily involves the stomach, in addition to the jaw, we reasoned the fictive feeding observed with optogenetic activation of CeA^Isl1+^ neurons may recapitulate the modulation of gastric function observed during normal eating. Increases in gastric acid secretion are a key aspect of gastric responses to the consumption of food. We accordingly predict that CeA^Isl1+^ activity may modulate gastric acid secretion during the consumption of food. To test this prediction, we first examined if gastric acid secreting cells have a poly-synaptic connection to CeA^Isl1+^ neurons (Fig. 5A). In GLP1r^Cre^ animals, we injected the Cre dependent poly-synaptic retrograde virus, PRV Introvert, into the gastric corpus. Four to five days after gastric injection of this virus, animals were perfused for conventional immunofluorescence on thin sections. Most (93.5%) of the PRV+ cells in the central amygdala stained positive for Isl1+, suggesting a specific connection between gastric acid secreting cells in the stomach, and CeA^Isl1+^ neurons.

**Figure 5:**
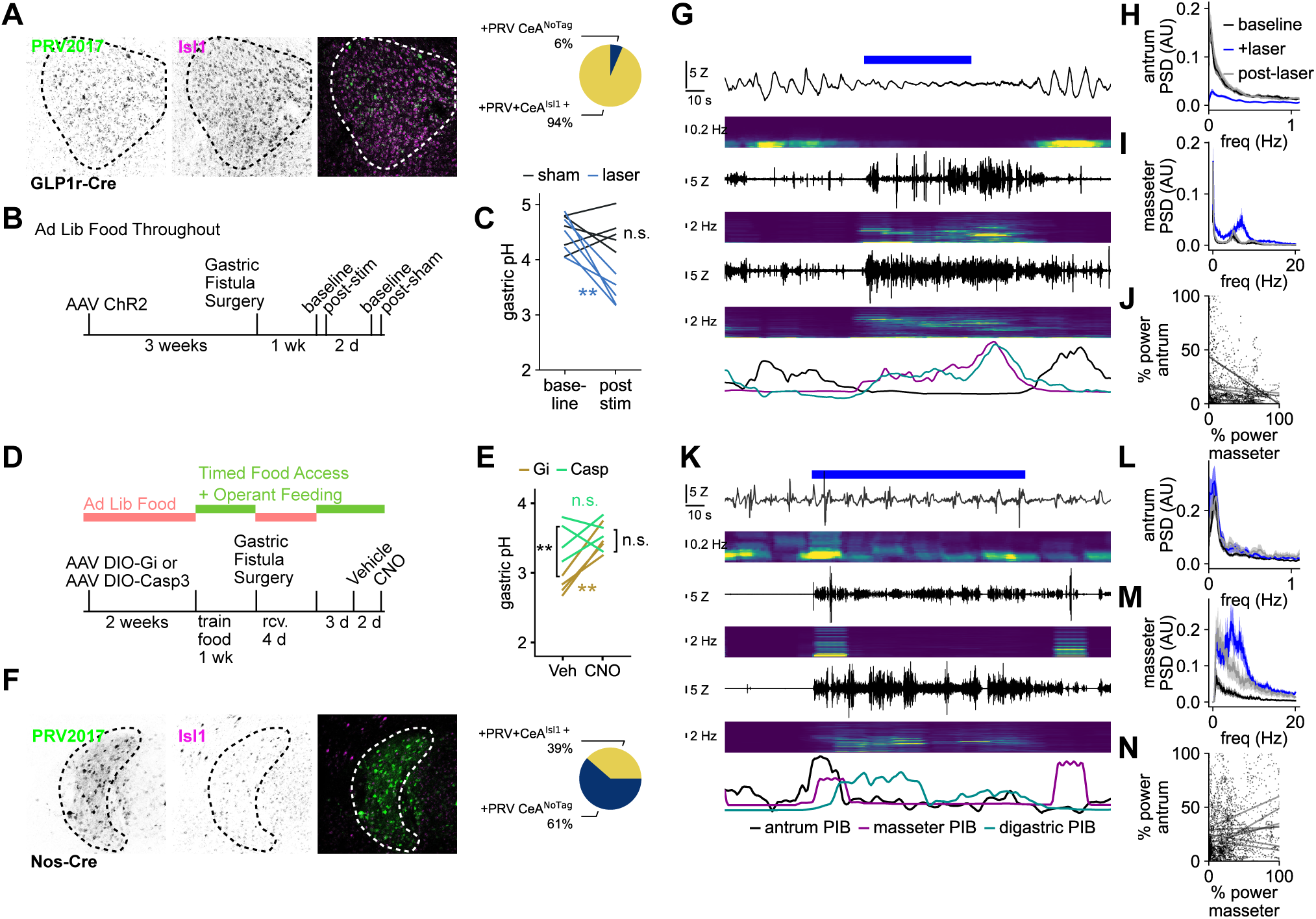
CeAIsl1^+^ neurons modulate gastric function. A-C. CeA^Isl1+^ are synaptically connected to gastric acid releasing cells in the stomach, and stimulating them reduces gastric pH. A. The Cre dependent poly-synaptic retrograde virus PRV2017 was injected in the gastric corpus in GLP1r-Cre mice. A substantial majority of PRV labeled cells in the central amygdala are positive for Isl1. B. Paradigm of experiment to observe effect of CeA^Isl1+^ stimulation on gastric pH (N=5). C. Activation of CeA^Isl1+^ cells reduces gastric pH. Laser activation reduced gastric pH (t_(4)_=5.65, P=0.005), whereas sham stimulation had no effect (t_(4)_=-0.031, P = 0.98) D-E. Activity of CeA^Isl1+^ neurons is necessary to reduce gastric pH in anticipation of a meal. D. Paradigm of experiment to observe whether inhibition or ablation of CeA^Isl1+^ neurons interferes with anticipatory gastric acid secretion E. Inhibition or ablation of CeA^Isl1+^ cells prevents reduction in gastric pH during meal anticipation. CNO treatment increased gastric pH in Isl1^Cre^ mice expressing the inhibitory DREADD in the CeA (t_(3)_=7.66, P=0.005), whereas CNO had no effect on animals receiving the caspase lesion of CeA^Isl1+^ neurons (t_(3)_=-0.382, P=0.728). Moreover, following injection of CNO, the gastric pH of the lesion group was not different from that of the Gi group (t_(6)_=0.768, P = 0.472), whereas after vehicle the gastric pH of the Gi group was significantly less than that of the lesioned group (t_(6)_=4.434, P=0.004). F-M. CeA^Isl1+^ are synaptically connected to enteric nervous system inhibitory neurons, stimulating them reduces gastric motility, and this effect depends on the integrity of the vagus nerve. F. The Cre dependent poly-synaptic retrograde virus PRV2017 was injected in the gastric corpus in NOS1-Cre mice. A minority of PRV labeled cells in the central amygdala are positive for Isl1. G. Representative EMG recordings of gastric and jaw signal before, during, and after stimulation of CeA^Isl1+^ neurons, laser stimulation indicated by blue bar at top. Traces in spectrograms are in pairs: upper is gastric antrum, middle is masseter, lower is digastric. Note the frequency range for the gastric antrum spectrogram is much lower than for the masseter and the digastric. H. Power spectral density estimates of the gastric antral rhythm before (baseline), during and after laser stimulation. Stimulation of CeA^Isl1+^ neurons significantly depressed the amplitude of gastric antral rhythm (t_(5)_=4.067, P=0.0097), while evoking rhythmic activity in the masseter (I). J. A significant negative correlation between the amplitude of the power spectrum of the antral rhythm and the amplitude of the chewing rhythm in the masseter is observed in 4/6 animals. K-N. CeA^Isl1+^ stimulation in animals receiving a bilateral subdiaphragmatic vagotomy. K. Example traces as in G. L. Power spectrum as in H, note that in vagotomized animals stimulation has no effect, (F_(2,14)_=1.19, P=0.33). M. Power spectrum as in I, stimulation still increases masseter activity, (F_(2,14)_=7.09, P=0.0074), laser stimulation is greater than baseline (t_(7)_=-6.7, P=0.00027) N. None of vagotomized animals had a significant negative correlation between masseter and antral activity.

Considering this specific anatomical connection between CeA^Isl1+^ neurons and cells driving gastric acid secretion in the stomach, we tested whether stimulating CeA^Isl1+^ neurons was sufficient to change gastric acid secretion. To do this, we expressed ChR2 in the CeA of Isl1^Cre^ mice, and after recovery from stereotaxic surgery, we created a chronic fistula window into the stomach of these animals so that we could repeatedly measure gastric pH. After recovery from the fistula implant surgery, we measured the pH of the collected gastric contents before and then again after 30 minutes of either sham or laser stimulation (Fig. 5B-C). Food was removed from the cage 30 minutes prior to laser or sham stimulation, and for the duration of the stimulus. Activation of CeA^Isl1+^ reduces gastric pH. A repeated measures ANOVA comparison (laser X time) found a significant interaction, F_(1,4)_=9.48, P=0.037, N = 5 animals. Post-hoc repeated measure t-tests confirmed that laser stimulation significantly reduced gastric pH by −1.06±0.19 (t_(4)_=5.65, P=0.005). Sham stimulation had no effect t_(4)_=-0.031, P = 0.98.

We next sought to test whether the activity of CeA^Isl1+^ was necessary for the changes in gastric acid secretion observed in anticipation of a meal. We elected to focus on anticipatory acid secretion because consumption of food increased gastric pH. In a pilot experiment measuring gastric pH in intact animals, as a terminal experiment, we observed a statistically higher pH (4.39±0.37) in the group that had access to food as compared to the food deprived group 2.92±0.17, (t_(6)_=3.61,P=0.011, Supp. 5A). Further, measurement of gastric pH in animals instrumented with gastric fistulas showed an increase in gastric pH following consumption of food that took nearly 8 hours to return to the fasted pH level (Supp. 5B). To test whether CeA^Isl1+^ neuron activity was necessary for anticipatory gastric pH changes, we prepared two groups of Isl1^Cre^ animals with chronic gastric fistulas. One group was an ablation group, where CeA^Isl1+^ are lesioned by injecting an AAV driving Cre dependent expression of Casp3. The second group was an inhibition group, receiving an injection of an AAV driving Cre dependent expression of hM4Di. Following recovery for stereotaxic surgery, animals are moved to a timed food access schedule and trained in an operant poking for food task, at an FR1 and then FR5 schedule. After acquiring the food poke task, gastric fistula surgery is performed. During this time the animals are returned to adlib food access to facilitate health and recovery. After 4 days of recovery, the animals are retrained in the food poke task for 3 days, they returned to timed food access, and poking for food in the operant box once per day. On the fourth day after retraining animals are dosed with CNO (or vehicle) and placed in the operant box to poke, but no food is provided, gastric pH is measured 10 minutes after they have been in the operant box. Two days after this subjects are crossed to the opposite treatment, i.e. vehicle (or CNO), placed in the box for 10 minutes to poke without food and the pH of their gastric contents is measured (Fig. 5D-E).

Inhibition or ablation of CeA^Isl1+^ cells increases gastric pH during meal anticipation. A repeated measures mixed ANOVA (Virus X Injection) found a significant interaction, and post-hoc paired t-tests confirmed that CNO treatment increased gastric pH from 2.81±0.62 to 3.46±0.10 in animals expressing the inhibitory DREADD (t_(3)_=7.66, P=0.005, Fig. 5E). In contrast, CNO had no effect on animals receiving the caspase lesion of CeA^Isl1+^ neurons, gastric pH after vehicle injection was 3.50±0.14 not significantly different than the gastric pH after CNO injection 3.58±0.11 (t_(3)_=-0.382, P=0.728). Moreover, following injection of CNO, the gastric pH of the lesion group was not different from that of the hM4Di group (t_(6)_=0.768, P = 0.472), but following the injection of vehicle, the gastric pH of the hM4Di group was significantly less than that of the lesioned group (t_(6)_=4.434, P=0.004).

### CeA^Isl1+^ cells modulate gastric motility

In addition to changes in gastric acid secretion, gastric motility also changes during consumption of food. Based on the preceding result, showing the involvement of CeA^Isl1+^ neurons on gastric acid secretion, we predicted that these cells may also be involved in modulating gastric motility. Both increases and decreases in gastric motility have been reported with consumption of food. Because we observed a decrease in gastric motility during consumption of food (Supp. 5A-D), we focused on circuits that could reduce motility. First, we examined if cells that slow gastric motility have a poly synaptic connection to CeA^Isl1+^ neurons. In Nos^Cre^ animals we injected the PRV Introvert in the gastric antrum and corpus. A minority of PRV positive cells (39%) were positive for Isl1, suggesting that it may be possible for CeA^Isl1+^ neurons to alter gastric motility by modulating the activity of gastric NOS neurons (Fig. 5F).

We next tested whether stimulating CeA^Isl1+^ neurons was sufficient to change gastric motility. A group of Isl1^Cre^ animals was injected with a Cre dependent AAV driving ChR2 expression in the CeA (N=6). After recovery from the stereotaxic surgery, wires were implanted to record electromyographic activity from the gastric antrum, as well as from the masseter and digastric muscles (Fig. 5G-J). In some animals duodenal EMG activity was recorded as well (N=3). Gastric antral EMG recordings showed a characteristic high amplitude, low frequency rhythm around 0.1 - 0.2 Hz, duodenal rhythm was characteristically distinct (Supp. 5I-L). Stimulation of CeA^Isl1+^ neurons suppressed the amplitude of this antral rhythm. During laser stimulation of CeA^Isl1+^ neurons, the amplitude of the power spectrum between 0 and 0.3 Hz is significantly changed (F_(2,10)_=15.1, P=0.00095), post-hoc tests show a reduction t_(5)_=4.067, P=0.0097 during laser stimulation as compared to the baseline period immediately preceding laser stimulation. A similar change to the antral activity was observed during the consumption of food, which significantly suppressed the amplitude of the antral power spectrum (F_(2,8)_=25.18, P=0.00036, Supp. 5A-C). Post-hoc tests show a reduction (t[4]=7.06, P=0.0021) during food consumption as compared to the baseline period immediately preceding consumption. The effect to reduce the gastric EMG appeared to be specific, as the duodenal rhythm was no significantly altered by stimulation (Supp. 5I-L). As observed previously, this same stimulation activated the masseter, increasing the amplitude of its power spectrum between 2 and 8 Hz (t_(5)_=-3.54, P=0.017) as compared to baseline.

We examined whether this apparent inverse effect of laser stimulation on antral and masseter activity exhibited a statistically significant correlation in our recordings. Segments of data recorded during laser stimulation were analyzed using time varying frequency estimates. Power spectra were computed with a sliding twenty second window moved across the data in 1 second increments. The total power of the relevant frequency band was computed for each muscle at each time point. The spectral power of the antrum signal was plotted as a function of the spectral power of the masseter signal, and linear regression analysis was performed. A significant negative correlation was observed between masseteric activity and antral activity for every animal recorded (Fig. 5J). A similar negative correlation was observed between masseter activity and antral activity during consumption of chow (Supp. 5D).

Having observed a negative correlation between masseter activity and gastric activity, we were curious to see if the effect to reduce gastric contractions was related to the chewing behavior elicited by CeA^Isl1+^ neuron stimulation, or a consequence of CeA^Isl1+^ activation *per se*. We reasoned that direct stimulation of the CeA^Isl1+^ terminals that project to the NTS would be likely to modulate gastric contractions, possibly without evoking rhythmic chewing or licking behaviors. To test whether CeA^Isl1+^ neuron activation reduced gastric contraction without evoking chewing behaviors, we injected a Cre dependent AAV driving ChR2 expression in the CeA, and placed optical fibers over the terminal projections in the NTS. After recovery from stereotaxic surgery, animals were implanted with electromyographic wires to observe changes activity in the masseter, digastric, gastric antrum (stomach) and duodenum, (N=5).

Laser stimulation of CeA^Isl1+^ terminals was less effective at evoking rhythmic oromotor behaviors. With the optical fiber placed above the central amygdala, masseter activity in response to 6 Hz laser activation is significantly greater than the pre-stimulation baseline, (Power in band 4–8 Hz, F_(2,8)_=11.97, t_(4)_=-7.67,P=0.0015). This was not the case when the fibers were placed above the NTS, stimulation of CeA^Isl1+^ NTS terminals at the same rate did not significantly change masseter activity (Power in band 4–8 Hz, F_(2,8)_=0.27,P=0.81). Nevertheless, activation of CeA^Isl1+^ NTS terminals at this rate significantly changed gastric activity (Power in band 0.05–0.2 Hz, F_(2,8)_=5.13, P=0.037). Post hoc comparisons showed that gastric activity was significantly increased during laser stimulation, as compared to the preceding baseline (t_(4)_=-3.47,P=0.0257). Higher frequency stimulation of the CeA^Isl1+^ NTS terminals, at 30 Hz laser activation also significantly changed gastric activity (F_(2,8)_=8.094,P=0.012). Post hoc tests showed a trend to reduce gastric activity during the laser stimulation (t_(4)_=2.48,P=0.068) and a significant rebound increase in gastric activity as laser stimulation halted (t_(4)_=-4.17,P=0.014). In contrast to this effect to potentiate gastric activity with laser activation of NTS terminals, stimulation of CeA^Isl1+^ neuron cell bodies in the CeA at the same rate significantly reduced gastric activity (F_(2,8)_=17.11,P=0.0013, t_(4)_=4.44,P=0.0113).

Inhibition of gastric motility is canonically associated with the actions of the sympathetic nervous system, activation of which can also reduce gastric acid secretion. Because CeA^Isl1+^ stimulation appears to increase gastric acid secretion, while reducing gastric motility, we reasoned that the effect on motility is mediated by the parasympathetic system rather than the sympathetic system. If the parasympathetic is involved, then cutting the vagus nerve should eliminate the motility reducing effect of CeA^Isl1+^ stimulation. To test this prediction, Isl1^Cre^ animals expressing ChR2 in the CeA were instrumented to record EMG activity in the masseter, digastric, and gastric antrum (N=7). During the EMG implant surgery, a bilateral resection of the subdiaphragmatic vagal nerves was preformed (Fig. 5K-N). Following recovery from surgery, we observed that CeA^Isl1+^ stimulation has no significant effect on the gastric rhythm (F_(2,14)_=1.19, P=0.33), despite still having a clear effect to increase masseter activity, (F_(2,14)_=7.09, P=0.0074). Post-hoc tests show that similar to intact animals, the amplitude of the masseter power spectrum between 3 and 8 Hz is increased by laser stimulation (t_(7)_=-6.7, P=0.00027). Further, The negative correlation between the masseteric activity and antral activity was eliminated. In vagotomized animals, during epochs of CeA^Isl1+^ optogenetic stimulation, only three out of seven animals showed a significant correlation between the masseter and the antrum, and for each it was a positive rather than negative correlation. In addition, in vagotomized animals (N=4) food consumption did not have the same effect on antral activity as observed in intact individuals (Supp. 5E-G). Instead, a significant increase was observed immediately following consumption (F(2,y)=6.9, P=0.028, t_(3)_=-4.3, P = 0.023). Moreover, none of vagotomized animals had a significant negative correlation between masseter and antral activity (Supp. 5H). After concluding these recordings, completeness of the vagotomy was assessed by examining anatomical spread of the retrograde tracer, ctb (Supp. 5M).

## Discussion

Animals engage in ingestive behaviors^5,7,8^ loosely referred to as ‘fictive feeding’ with stimulation of the central amygdala. Here we examine in some detail the extent to which ‘fictive feeding’ produced by optogenetic stimulation of CeA^Isl1+^ neurons recapitulates the entire suite of physiological changes observed during ingestion. In addition to biting and chewing activities, we also observe changes in gastric acid secretion and gastric motility without the ingestion of food. Activation of a single molecularly-defined population of neurons is capable of eliciting and coordinating both gastric and oromotor behaviors of ingestion.

### Generation of High Force Bites

Stimulation and inhibition experiments presented in Figure 1, indicate that CeA^Isl1+^ neurons are sufficient and necessary for the generation of high force bites. In stimulation experiments, both the frequency of bite generation and the force of each bite showed a significant positive relationship to the rate of laser stimulation, suggesting these cells may support both the initiation of jaw movement and the generation of force. Inhibition of CeA^Isl1+^ neurons via chemogenetic means had an effect to diminish both the generation of force, and the initiation of biting. Both movement initiation and force generation are linked to motivational processes, suggesting a third function for CeA^Isl1+^ neurons, to establish a motivational state. During the CeA^Isl1+^ stimulation experiment, the motivation to bite the force meter is unclear. In experiments where the force meter was replaced with a dry sipper, during lower frequency stimulation, animals would lick the sipper, suggesting a cue dependent appetitive component to the motivational state established by CeA^Isl1+^ stimulation. Regardless, at high rates of stimulation, the predominant motor behavior evoked was biting, and the number of bites evoked when either the force meter or a dry sipper was presented as a target, increased with increasing rate of stimulation. We suggest that the activity of these cells may be intrinsically linked to the behavior of biting, and this association seems to supersede licking.

Recordings from CeA^Isl1+^ neurons during an operant task showed that a portion of these neurons have significant firing rate correlations with bite force. These correlations existed over a short time frame, in the 200 millisecond around individual bites. In addition to neurons significantly correlated with bite force, a proportion of CeA^Isl1+^ neurons were specifically active during licking. Cells that were significantly active during the generation of individual bites were typically not significantly active during the generation of individual licks. Many CeA^Isl1+^ neurons that exhibited elevated rates of firing when the animal bit the force meter were also highly active during the consumption of food. These data suggest that the CeA^Isl1+^ population is not functionally homogeneous, and that individual cells may support licking or biting, regardless of the sensory target.

### Involvement of the Periodontal Masseteric Reflex

Supporting the idea of an intrinsic link between CeA^Isl1+^ neurons and biting, we describe a specific projection from CeA^Isl1+^ neurons to Me5 sensory neurons that project to the periodontal ligament. In cats and rats, a similar anatomical connection has been previously described between the mesencephalic trigeminal ganglia and the central amygdala^9–11^. The projection of the peripheral endings of mesencephalic trigeminal sensory neurons into the periodontal ligament is specifically associated with the incisor teeth. In rats, cats, and dogs Me5 sensory neurons invest the periodontal ligament on the ab-lingual face of the incisor, or canine teeth^12^. The periodontal ligament receives dual innervation from neurons having their cell bodies in either the trigeminal ganglia or the mesencephalic trigeminal ganglia. These two populations of sensory neurons have different distributions of their terminal sensory endings within the periodontal ligament. Trigeminal ganglia sensory neurons do not show the preference for the ab-lingual face of the canine in cats that is seen for Me5 neurons. A substantially higher proportion of mesencephalic trigeminal neurons are responsive to forces on the tooth in the linguo-labial direction, as compared to cells found in the trigeminal ganglia^13^. During biting, the canine teeth would experience forces in the linguo-labio direction.

Tapping rat teeth in this linguo-labio direction causes a reflex contraction of the jaw closing muscle, an effect that cat be potentiated by direct electrical stimulation to the mesencephalic trigeminal nucleus^14^. In the cat, electrical stimulation of the Me5 causes a similar reflex contraction of the masseter, and this effect can be potentiated by stimulation to portions of the amygdala^15^. The anatomical and physiological projections from CeA^Isl1+^ neurons to Me5 neurons presented here suggest a similar circuit is present in mice. The synaptic projection from CeA^Isl1+^ neurons appears to be depolarizing projection, sensitive to blockers of GABA_A_ receptors. Depolarizing responses to GABA have been previously in Me5 neurons^16^ and the reversal potential measured here is similar to other estimates^17^. Further it seems that the depolarizing effect of CeA^Isl1+^ stimulation, which occasionally caused Me5 neurons to spike, had an excitatory and facilitative effect. This is in contrast to the shunting effect on spike propagation that has been described with stimulation of brainstem regions adjacent such as the lateral supratrigeminal^18^. Taken together, these results fit a model where CeA^Isl1+^ neuron activation facilitates Me5 excitation of jaw-closing motor neurons in response to sensory stimulation to the teeth. Supporting this hypothesized role, we observed that the effect of lesioning Me5 neurons is to reduce bite force, without reducing the number of bites generated.

### Effects of CeA^Isl1+^ neurons on motivation

Reports of brain regions that can support self-stimulation behavior frequently find an inherent link with food consumption. Self stimulation of midbrain dopamine neurons is facilitated by food deprivation, and self-stimulation vigor is reduced by re-feeding^19^. Lateral hypothalamic self stimulation reduces food intake, and force feeding or gastric distension can increase current thresholds for self stimulation in this region^20^. Here we report a similar finding, that CeA^Isl1+^ neurons which can function as an operant reinforcer to support intra-cranial self stimulation are linked to the ingestion of food. Interestingly, we do not see the same effect observed with lateral hypothalamic self stimulation. Food restriction does not seem to change the frequency with which animals self-stimulate. This points to a central motivational difference between the lateral hypothalamus and the CeA^Isl1+^ neurons described here. In response to stimulation of CeA^Isl1+^ neurons, hungry animals will not orient to food that is out of their visual field. In contrast stimulation of lateral hypothalamic areas can facilitate ingestive behavior that involves orienting to and consuming food in preference to other non-nutritive materials^21^. Further, inhibition of CeA^Isl1+^ neurons does not reduced food intake, unlike the effect seen with inhibition of the lateral hypothalamus in rats^22^. Despite the fact that inhibition of CeA^Isl1+^ neurons had no effect to reduce food intake, it did seem to interfere with animals ability to consume food with their normal facility. In particular, chemogenetic inhibition of CeA^Isl1+^ neurons caused animals to shred significantly more of their food.

### Contributions of CeA^Isl1+^ neurons to gastric functions

A wide variety of manipulations of central amygdala activity have been shown to modulate gastric function. Electrical stimulation of specifically the medial portion of the nucleus has been shown to increase motility, whereas stimulus to the central lateral amygdala has the opposite effect^23^. Microinfusions of TRH and orexin both increase gastric contractions^24,25^, and TRH infusions can increase gastric acid secretion^26^. Infusions of nesfatin-1, a gut hormone that can reduce gastric acid secretion^27^, into the CeA reduces gastric motility^28^. Stimulation of CeA^Isl1+^ neurons had an effect to reduce gastric pH, and inhibition of these neurons in animals anticipating a meal increased it. This result suggested that CeA^Isl1+^ neurons may play a role in the cephalic phase of ingestion, specifically on the type of anticipatory gastric secretions first described by Pavlov^29^. Inhibition experiments suggest that CeA^Isl1+^ neurons maybe active in animals anticipating a meal. If CeA^Isl1+^ neuron activity supports biting, we predict that gnawing or biting would occur more frequently in mice anticipating a meal. Unfortunately, we did not record jaw EMGs during this experiment, and are unable to find reports of increased oromotor activity during food anticipation. How these neurons’ activity changes during the transition from anticipatory behaviors to consummatory behaviors is an interesting question we do not address here.

The effect of stimulating CeA^Isl1+^ neurons on gastric motility is less clear. The predominant effect on motility we observed was a transient reduction in gastric contractions. In one circumstance we observed an increase in gastric contractions, when stimulating CeA^Isl1+^ terminals in the NTS, using rates of laser activation that did not elicit oromotor behaviors. Based on these conflicting findings, it maybe that different projections of CeA^Isl1+^ neurons could have opposing effects on gastric motility. Specifically, the NTS projections of CeA^Isl1+^ neurons appears to target the ventral-commissural NTS, which contains a populations of gastric stretch sensitive neurons. Electrical stimulation of the vcNTS reduces gastric motility. The neurons in the vcNTS extend dendrites ventrally to the parvocellular reticular area^30^. Chewing related activity in the parvocellular reticular area could inhibit gastric activity, possibly by exciting these neurons. Chewing gum after consuming a liquid meal can slow its initial rate of emptying from the stomach^31^, indirectly suggesting that chewing can inhibit gastric motility. One report describing observations made during surgery recounted a dramatic inhibition of the typical ongoing gastric contractions when the surgical patient was suggested to chew gum^32^. Inhibition of the ventral commissural NTS neurons by CeA^Isl1+^ projections would be expected to increase motility.

### Summary

The performance of specific ingestive behaviors involves coordinating the activity of the stomach and the jaw. We describe here a population of neurons that seems to be critical for the generation of forceful bites, and is active during consumption of food. This same population also lowers gastric pH during meal anticipation, and can transiently suppress gastric contractions. These cells seem to facilitate the consumption of food, at both the level of conscious consumption, and unconscious digestion.

### Support

This work was performed with the support of the NIH: R01AT011697

## Methods

### Measurement of Bite Force

The bite force of mice was measured using a custom fabricated bite force meter. This device has been described previously,^6^. A purpose-built mouthpiece, whose dimensions (H = 1mm × W = 5mm) were based on incisor morphology of adult C57BL6/J mice, was affixed to an accurate single point load cell system (OEM Single Point Load Cell LCAE-3KG), coupled to a signal conditioner (IN-UVI Omega). To reduce 60 Hz hum, which lead to inaccurate peak detection in the force profiles, it was necessary to power the signal conditioner with a precision linear regulated DC supply (Acopian A24MT210). The two bite plates of the force meter were spaced apart by 1.5 - 2 mm. Output signals were digitized and synchronized with other recorded signals (i.e. camera frames, muscle EMG, or single unit recording when performed) via a Tucker-Davis-Technologies RZ5. Awake animals were restrained for the biting tests, as is usually performed. Raw signals were low pass filtered (100 Hz) and peaks were detected for statistical analysis. Our baseline measurements in control mice (∼6-8N) are slightly less that published values of maximum bite force for healthy adult mice. In this study, we elected to place the bite plates relatively close, such that animals were not making a very large gape. Based on relationship between vertical jaw opening and maximal occlusal force^33^, this should reduce the maximal forces measured.

### Measurement of Bite Force During Isl1 Stimulation

Isl-Cre animals were injected with DIO-AAV-ChR2 (150 nL bilaterally in the central amygdala) and in the same procedure bilateral optical fibers were placed above the injection site for optical activation of ChR2. Following 3 weeks of recovery, animals were habituated to brief periods (around 60 seconds) of restraint in front of the bite meter, while connected to the light source. On the next day, the force of bites evoked by laser stimulation of Isl1 cells was measured. Mice were restrained with their snouts in front of the force meter, then a train of pulses, 8 seconds in duration, with a frequency of 2–40 Hz was given while the animal was in front of the bite meter. Trains of stimulation were presented in pseudorandom order with respect to frequency. The average force of each bite, the total impulse, and the number of bites was calculated for each trial. 2–3 trials of each frequency were performed for each mouse, and trials of the same frequency were averaged such that each mouse is represented by a single point at each stimulation frequency. In a subset of animals, EMG recordings were made by during the measurement of bite force. For each recording muscle, a pair of nichrome wires (A-M systems 761500) was stripped of insulation along a 0.5 mm segment and twisted together. Strands in the wire pair were arranged such that the bare regions were separated by about 1 mm once twisted together. The pairs of wires were crimped onto 30 gauge needles which were used to pass them into either the masseter or digastric muscle. Wires were solder trapped in into millmax connectors (832-10-006-10-005000) which were affixed by dental cement to the headcap.

### Lesioning of Me5 neurons

Isl1 is expressed in cranial nerve sensory neurons, including the mesencephalic trigeminal sensory neurons whose cell bodies reside in dorsal pons. Expression of the apoptotic enzyme caspase 3 was driven selectively in these Me5 sensory neurons by injection of AAV1 which caused Cre expressing cells to produce constitutively active Caspase 3, leading to their apoptosis. Because these neurons are the only cells in the parabrachial region expressing Isl1/Cre, and because AAV1 shows very little retrograde activity, this manipulation is expected to selectively eliminate the Me5 neurons. The more caudal portion of the Me5 region was targeted as this area appears to harbor relatively more of Me5 neurons projecting to the periodontal ligament34. In the same surgery AAV5-EF1a-DIO-hChR2(H134R) was injected bilaterally into the central amygdala, and optical fibers were placed about the central amygdala. Control animals of the same genotype (Isl1^Cre^) received AAV1-hSyn-DIO-hM4D(Gi)-mCherry injections in the place of AAV1-FLEX-taCasp3-TEVp.

### Food Intake Chow

Food intake was measured in the home cage for animals that had ad-libitum access to 20 milligram pellets of food (Bio-serv precision diet 20 milligram pellets) provided by a FED3 feeder. Animals were acclimated to FED3 feeder for 10 days, during this period their food intake became regular, with a noticeable increase in pellet retrieval starting about 2 hours after lights off. Starting at day 10, animals were habituated to brief restraint and injection of saline, 2 hours after lights off. After two days of saline injection, animals received a single dose of CNO on day 3, again at 2 hours after lights off. Following this injection on day 4, animals received saline at 2 hours after lights off. Timestamps from the FED3 for pellet retrieval were used to measure food intake in the 2 hours after injections. Two groups were compared, animals expressing ChR2 in the CeA, or animals expression HM4(Gi)-mCherry in the CeA.

### Food Intake Pasta

Isl1^Cre^ animals were first injected with either hChR2 or stGtACR2 in CeA, and fibers were placed above the CeA. 3 weeks after this surgery, animals were introduced to dry pasta (Barilla spagetti, cut into 2 cm lengths) in their home cage while they had access to standard chow (Lab diet 5053) for three days. Additionally, during this period, animals were taken individually from their home cage and placed in a round plexi-glass behavioral area to habituate them to behaving while connected to the fiber optic cables and commutator. On day 4 animals were food restricted for 16 hours. They were then placed in the circular arena, and given two 2 cm segment of pasta, sequentially (they got the second after finishing the first). Activation of 473 nm DPSSL laser was controlled manually by the experimenter, such that when the animal retrieved and brought the pasta to their mouth, the laser was activated to illuminate the CeA, and this was continued until the animal dropped the pasta. The pattern of laser illumination and amplitude was different between the two groups. For the GtACR2 group, the laser was continuously on when triggered, and laser amplitude was adjusted to be between 3-7 mW at the fiber tip. For the ChR2 group, the laser was pulsed at 10 Hz, with a pulse duration of 8 milliseconds, laser amplitude was adjusted such that it would evoke fictive feeding within 5 seconds of initiation about half of the time. Animals were randomly assigned to receive either laser activation during consumption of the first pasta piece, or during the second piece. The latency to finish each piece and the number of drops and re-initiations was counted.

Measurement of Jaw Muscle EMGs during pasta eating for GtACR2 animals. Following measurement of pasta eating as described above, animals from this group were implanted with EMG wires for the measurement of jaw muscle activity during pasta eating using the TDT-Medusa low impedance differential amplifier coupled to a TDT-RZ5. Electrode wires were fabricated as described in fabrication of EMG assemblies section.

### Measurement of shredded chow

Animals were singly housed and food restricted to 90% of their free-feeding body weight at the start of the experiment. In the early evening, animals were moved from their home cage to a plastic container. A total of 5.5 grams of chow (LabDiet 5053) were added to the container with the mouse. Two to three intact pellets were selected who’s weight added to the desired amount, when necessary some pellets were shaved with sharp razor to reduce their weight. Animals fed freely for one hour, at which time they were removed. After removing any droppings from the container, the remaining food was gentle sifted through a 1/8″ stainless steel sieve, (ADVAN-TECH 1/8″B Supp. 8H). The weight of the food passing the sieve, and that retained by the sieve were measured. The sum of these two weights, less the amount of food given at the start of the session was computed as the weight of chow consumed. The ratio of the weight of the food passing the sieve to that retained was used to compute the percentage of chow shredded. About half of the mice urinated during the food intake measurement. After removing the mouse, but before weighing or passing food through sieve, urine was dried by placing the plastic container in the hood, then the sieving and weighing were performed.

### Measurement of the periodontal-masseteric reflex

Isl1^Cre^ animals were injected with AAV5-ChR2 in the CeM and Fiber were placed above the CeA. 3 weeks later, animals were anesthetized for placement of EMG wires in the masseter and digastric muscles. Wires were fabricated as described in the EMG assembly section. Following implantation of EMG wires, animals recovered for at least 2 days. Next, on the day of the experiment animals were anesthetized with isoflurane and a titanium head fixation plate was added to the head cap, along-side the EMG connector and the fiber optic cannula, as cement was setting on the headplate, the mouse was transitioned to urethane anesthesia, 1.1 – 1.4 g/ kg. Isoflurane anesthesia was withdrawn while monitoring anesthetic depth. The animal was transferred to the recording area, headfixed, and EMG wires were connected to the medusa amplified (TDT RA16LDI). A camera (Basler aCA1300gm) was positioned so that responses could be visualized at a higher frame rate (170 fps). Frame strobes were recorded by the TDT for post recording synchronization. Video files were encoded using a gstreamer plugin for pylon cameras and Nvidia hardware h264 encoding. Once stable under urethane anesthesia, the EMG responses to manual stimulation of the oral cavity and teeth were recorded. Using a small blunt needle holder, the upper incisors were carefully pressed in either the labio-lingual direction or the linguo-labial direction. Responses to other stimuli were explored: gentle stroking of the upper lips, and a firm upward press on the hard pallet. Timing of the tooth stimulation was determined after completing the experiment by frame-by-frame examination of the video record. Frame accurate timestamps of tooth contact were extracted by taking millisecond timestamped screenshots using mpv. Time stamps were translated to recording times, and peristimulus EMG activity could be computed. EMG traces were rectified, integrated (convolution with a 10 millisecond boxcar), and then lowpass filtered at 200 Hz. This integrated EMG signal was used to generate peristimulus muscle amplitudes for figures. Animals were euthanized at the completion of the recording (perfused for histology).

### Simultaneous measurement of jaw muscle activity and gastric muscle activity

To examine coordination of muscle activity between the jaw and stomach, simultaneous EMG recordings were made of both structures. Isl-cre animals were first injected with AAV5 ChR2 and optical fibers were placed above the injection site. 3 weeks after this surgery, animals were prepared for surgery. A series of small patches were shaved, the upper neck below the headcap, between the scapula, and the ventral midline. First the recording wires were tunneled subcutaneously from the headcap to the scapula, then from there to the skin below the lowest rib. At this point, a ventral midline incision was made to expose the stomach and duodenum, and the recording wire were passed through the abdominal muscle by create a small perforation in the muscle about 1.5 cm lateral to the midline. The stomach and proximal duodenum were bathed in saline with 100 micromolar nicardipine, to reduce their movement and facilitate placement of the recording wires. Initial experiments employed home-made assemblies of wires as described in the Fabrication of EMG assemblies below. A small deviation from the approach used for skeletal muscle was employed, instead of using the standard twisted pair, 4-6 wires were arranged in parallel for a single recording site. This was done to allow signals to be referenced if one strand is broken, or the assembly moves. A second change was made, to reduce the frustrating breakage of entire wire assemblies, a length of 8-0 suture was incorporated into the wire bundles going to the gastro-intestinal sites. The suture and bundled wires were twisted together, and crimped into a 30 gauge needle. The needle carrying the wire bundle was carefully passed tangential to the stomach serosa, making an attempt to have it pass through the muscular layer as much as possible. Wires were placed so that for recordings from the gastric antrum, the bare recording sites of the wire bundle would rest between 2-5 mm from the pyloric ring. Once placed, the needle was clipped off, and the trialing wires were sutured to the serosa with fine 9-0 suture. In some experiments a second bundle was placed in the duodenum about 5 mm distal to the pylorus. After placement in the abdomen, the midline incision was closed, and next wires were implanted in the masseter and digastric. Simple twisted pairs were used for these recordings, the wires were routed subcutaneously behind the ear and then into the masseter and digastric.

### Myomatrix recording of gut and jaw muscle activity

Because of the difficulty in recording muscle activity in the gut and jaw simultaneously, we moved to a precision-engineered recording assembly, focusing on flexible tissue. Myomatrix Arrays were received from the CAMBER institute in Georgia. Arrays with 4 threads were used to target the gastric antrum, masseter, digastric, and duodenum. Surgical details are similar as for the placement of nichrome bundles described above. One modification was create a lasso that facilitated the placement of the thread along the muscular tissue in the stomach and duodenum. A 5-8 centimeter length of 8-0 nylon suture was passed through the last small hole at the tip of the myomatrix array, and folded in half. The two ends of the suture were carefully threaded into the blunted end of 30 or 31 gauge needle, which were then trapped in the needle by crimping the blunt end. This created a needle lasso that could be used to perforate the gastric or duodenal tissue and pull the myomatrix thread through without having the drag of a knot.

### Fabrication of multiple target EMG recording assemblies

Lengths of nichrome wire (A-M systems 761500) were bared of insulation in a small region (about 0.5 mm) about 5 cm from the end of wire to create a recording zone. Pairs of wires were arrange in parallel with their recording zones separated by about 1 mm. Holding the paired wires in an impromptu modeling clay clamp, they were passed through a short (0.5 mm) round of polyolefin primed(Premabond POP) silicon tubing (Braintree RenaSil Silicone Tubing .025 OD x .012 ID). The silicon tubing was positioned about 1.5 mm proximal to the bared recording zones on the wires and affixed by carefully filling with loctite 401, which easily wicked into the primed tubing. Next, A 30G 1/2″ hypodermic needle (Exel) was broken from its hub, being careful to maintain the sharp end. The blunted end was ground on an Arkansas stone, and then the wire pair was inserted into it and the needle was crimped onto the nichrome wires. This created a pair of recording wires in the same style described by Kier Pearson, but more flexible, and better suited to recording from smaller muscles. The pair of wire opposite the crimped needle was stripped and solder trapped into a connector (either MillMax 853-13-006-10-003000 for use with the passive TDT-Medusa amplifier or Neuralynx EIB16 for use with TDT RA16PA amplifier and Intan headstages). For each recorded muscle, two pairs of wires were constructed. The portion of the wire pairs closest to the headcap was passed through a ∼ 7 mm length of silastic tubing and sealed with Kwik-Sil. This helped to prevent the animal from breaking wires where they passed from the headcap to under the skin. A ground connection was added, a short 1.5 cm length of PFA insulated braided stainless steel (A-M systems 790500) was stripped of insulation for 2 mm and routed subcutaneously to the back of the neck. The connector was affixed to the headcap with a mixture of dental cement and loctite super glue gel control.

### Slice electrophysiology

The coronal slices containing the parabrachial area (PBN) were prepared from Isl-cre mice, that had previously been infused with AAV5 driving ChR2 in a Cre dependent manner. Briefly, mice were anesthetized with isoflurane and perfused with slicing solution and then decapitated. Then brains were rapidly removed and immersed in cold (4°C) and oxygenated low sodium solution containing (mM): 92 NMDG, 2.5 KCl, 1.25 NaH2PO4, 30 NaHCO3, 20 HEPES, 25 glucose, 2 thiourea, 5 Na-ascorbate, 3 Na-pyruvate, 0.5 CaCl2·2H2O, and 10 MgSO4·7H2O, pH titrated to 7.3–7.4 with HCl and bubbled with 5% CO2 and 95% O2. After being trimmed to a small tissue block containing the PBN, coronal slices (250 μm thick) were cut on a vibratome (Leica VT1200S) and moved to a recovery chamber at 30°C, and sodium was slowly reintroduced to the sections, by spike-in method. A bubbler chimney improved washing of sections during this recovery period. After 25 – 45 minutes of recovery at 30°C, slices were moved to a holding chamber at room temperature with HEPES aCSF (in mM): 92 NaCl, 2.5 KCl, 1.25 NaH2PO4, 30 NaHCO3, 20 HEPES, 25 glucose, 2 thiourea, 5 Na-ascorbate, 3 Napyruvate, 2 CaCl2·2H2O, and 2 MgSO4·7H2O. Titrated pH to 7.3–7.4 with NaOH, bubbled with 5% CO2 and 95% O2. After recovery at room temperature for at least one hour, slices were transferred to a recording chamber constantly perfused with aCSF at a temperature of 33°C and a perfusion rate of 2 ml/min for electrophysiological experiments. Whole-cell patch clamp recording was performed in large neurons lateral to the LC (putative Me5 neurons) under both voltage and current clamp. Me5 neurons were identified by their intrinsic electrical properties, namely a prominent sag current, and 2-5 mV oscillations in the frequency range between 150 – 200 Hz during prolonged depolarizing steps. Micropipettes (3-4 MΩ) were made of borosilicate glass (World Precision Instruments) with a Sutter P-97 micropipette puller and back filled with a pipette solution containing (mM): K-gluconate 110, KCl 25, MgCl2 2, HEPES 10, EGTA 1.1, Mg-ATP 2.5, Na2-GTP 0.3, and Na2-phosphocreatin 10, pH 7.3 with KOH35. Both input resistance and series resistance were monitored throughout the experiments and the former was partially compensated. Only recordings with stable series resistance and input resistance were accepted. To stimulate neurons with an optogenetic method, an LED-generated blue light pulses at different frequencies (5, 10 and 20 Hz) were applied to recorded neurons. All data were sampled at 10 kHz using a CED 1401 micro3. Analysis was performed using CED’s Spike2 software.

### Inferior Alveolar Nerve Injection

To identify Me5 neurons projecting to the periodontal ligament, 1% flourogold was injected into the inferior alveolar nerve. Isl1^Cre^ animals first were injected with AAV1-hSyn-FLEx-mGFP-2A-Synaptophysin-mRuby in the CeM. Four weeks after this injection, animals were prepared for surgery, an incision was made to expose the masseter muscle. Using two blunt probes the masseter muscle was separated to expose the bony prominence of the mouse mandible through which the IAN courses. To assist in exposing the nerve and injecting into it, a small clamp was used to hold the lower jaw in place while still providing anesthetic gas. Using a dental drill, a small window was created in the prominence. A beveled glass needle containing 1% fluorogold was carried on a manual manipulator (Prior) and inserted through this window. Using a picospitzer about 0.5 microliter of FG was injected into nerve canal. The window was sealed with bone wax and the muscle and skin sutured back into place. Animals were perfused for histology 7 days later.

### In vivo electrophysiological recordings with opto-tagging

First, Isl1-Cre mice were injected with AAV5-DIO-hChR2-xFP into the CeA on either the left or right side. 2-3 weeks following the virus injection, mice were placed on the stereotaxic apparatus, and a pulled/tapered optic fiber (200 micro 0.22 NA, 1 mm taper Doric) was placed at an 20 degree medial/lateral angle relative to plumb, such that the tip rested at the nominal atlas coordinate of AP: 1.0, ML: 2.6, DV: 4.5. The fiber was secured to an adjacent headscrew (00-90 1/16, 18-8 SS, torx T2, McMaster-Carr #90910A380) with loctite 401 and dental cement (SNAP). Next, a micro-drive containing a 16-channel bundle of individual tungsten wires (Innovative Neurophysiology, Inc. USA) was placed above the CeA (AP: 1.0 mm, ML: 2.5mm, DV: 3.5-3.8 mm). Two days after recovery from the virus surgery, the wire bundles were lowered 50 microns at a time while recording and pulsing the laser to search for opto-tagged units. The locations of electrodes were confirmed histologically. Recordings were performed using the spike modules of the multichannel acquisition processor (Tucker-Davis Technologies).

### Identifying opto-tagged units

In each opto-tag recording session, a sequence of 20 - 200 laser pulses were driven, with a pulse duration was between 20 and 200 milliseconds, and inter pulse interval was between 2 and 5 seconds. Laser amplitude was manually adjusted while monitoring by audio and oscilloscope. Starting from a low level, laser amplitude was increased until that the unit latency was minimal and then decreased so that the waveform was clearly discernible from any opto-electric artifact, and the audio signal gave an impression of repeated impulses not diminishing during the period of illumination. Once the amplitude was established, a sequence of pulses was given, and unit firing in this sequence was used in the stimulus associated spike latency test (SALT), indicating whether the cell could be considered to express ChR2, meaning it is a putative Isl1+ central amygdala neuron.

### Operant Bite Force Task (Tootsie-Pop task)

To determine whether Isl1+ central amygdala cells firing is related to bite force under conditions of voluntary biting in a situation that approximates a normal motivated behavior, we developed a novel operant task. The basic idea was to couple the application of a forceful incisor bite to the provision of a food reward, in a manner like that which could occur during ingestion, we call this the tootsie-pop task. The same bite force meter used to measure bite force during optogenetic stimulation, or during restraint was passed through a cut out in a med-associates rat behavior box (ENV-007-VP), so that it rested about 1 cm above the grid floor. An intra-oral cannula was surgically implanted into an Isl1^Cre^ mouse previously instrumented for opto-tag recording of Isl1+ units in the CeA. Before training, animals were food restricted, chow was removed from their cage 16 hours before the session. For the first session, Recess peanut butter filling was pressed in between the two bite plates of the force meter, so that animals are more likely to nibble on the force meter. Oral infusions of intra-lipid were done by pressure. The intra-oral catheter was connected to a reservoir of 20% intra-lipid which was under nitrogen pressure 1-5 psi. Flow from this reservoir into the mouse’s oral cavity was controlled by a normally closed solenoid (coleparmer 98302 12VDC/15psi). Because of the relatively high compliance of the tubing, our first attempt, using a syringe pump, introduced a substantial delay between the moment of biting and the receipt of the oral reward, which made learning the task in a single session difficult. Using a pressure delivery system, this latency was reduced, and animals could learn the task in a single session, and all reliably learned the task in 2 sessions. Valve open time was fixed at 200 milliseconds, and pressure was adjusted so that approximately 10 microliters of intralipid was delivered for each activation of the valve. The initial force threshold was set to 0.3 N, as mice could only generate this much force with their jaws, no animals were able to exceed this threshold by manipulating the force meter with their paws. Both unit activity and bite force were monitored using a TDT RZ5, which was also used for closed loop activation of the intra-lipid infusion solenoid. A small opto-isolator control circuit (BOB-09118) was used to drive a MOSFET (FQP30N06L) powering the solenoid, solenoid closure was passive (Schottky diode for flyback). In the task, the force-threshold for which a reward oral infusion was given incremented every 10 rewards earned. In the first and second session the amplitude of the threshold-increment was 0.1 N, in subsequent sessions it was 0.3 N, in addition, a speaker was linked to the force meter, such that for the duration of time the force meter was above threshold, a tone played. After the second session, the force meter was not baited with peanut butter, and the tone was eliminated, except for occasional manually activation to test whether recorded units responded to the tone.

### Recording of single unit activity during licking

In addition to recording Is1l neuron activity during biting, in a subset of animals performing this task, activity of neurons during licking was also recorded. To get precise timing of lick onsets and offsets, we preferred to use a contact based lick-o-meter, as optically based lickometers seemed difficult for the animals to access with their head caps. To avoid the electrical artifacts introduced by current-passing contact lickometers, we constructed a comparator based lickometer36. A custom grid floor was constructed for the Med-Associates box using aluminum rod instead of the typical stainless steel. The fluid sipper was a stainless steel. This created a metal potential difference that could be measured when the animal licked the stainless steel while standing on the aluminum grid flood. Lick onsets and offsets were compared to a manually adjusted threshold and converted into a digital level allowing TTL lick-on and lick-off events to be recorded using the same digitization hardware as was used for unit recording, a TDT RZ5.

### Intra-oral cannula placement

Animals were anesthetized and prepared for surgery as described above. A 2.5 cm segment of polyurethane tubing (micro-renathane OD: .033″, ID: .014″, Braintrain MRE033) was passed subcutaneously from the headcap to the portion of the cheek between the incisors and molar which is free of muscular. At the headcap, the catheter was passed though a short length of hypodermic tubing (deburred blunted 18 gauge needle) and secured to this conduit of tubing with a mixture of dental cement and loctite 401. Oral-catheter tubing could be connected using a blunted 26 gauge needle to an longer length of the same tubing for intra-oral infusions. Animals were left to recover for two days after placement of the oral catheter before performing any recording or behavioral tasks.

### Pseudorabies virus tracing

The abdomen of 8-hour food restricted animals was shaved and cleaned. As the animal was and prepared for surgery, a 5 mg/kg dose of dexamethasone was administered, this has been reported to facilitate alpha-herpes virus replication. A midline incision was made into the abdomen. Stomachs were exteriorized through the midline incision, into a bath of saline with 100 micro-molar nicardipine added to reduce smooth muscle contractions and facilitate needle placement in the myenteric layer of the gastric antrum. PRV-Introvent-GFP70 was mixed 5:1 (v:v) with 0.1% fast green in a hypertonic saline solution (500 mOsm), to facilitate visual observation of injection. Virus was front filled into filament free glass capillary (1.0 OD, 0.5 ID Sutter) which had been pulled on a Flaming-Brown puller (Sutter P-97), broken with a ceramic tile, and then beveled using home-made dry beveler. Briefly: diamond lapping (EMS 0.1 um grit) was adhered to a western digital hard-disk platter, using a manual manipulator, (Prior) the glass needle was lowered carefully onto the spinning disk platter and beveled for 30-40 seconds at a 30 degree angle, under positive pressure to prevent clogging. Pipette tips were visually inspected for a clean sharp appearance with a 10-20 micrometer diameter opening and a single faceted bevel. Glass needles loaded with virus/fast green mixture were connected to a picospritzer (MSI-200), and the needle tangentially inserted into the gastric antrum, such that with application of pressure, a vaguely discernible honeycomb-like tiling of fast-green tinted injectate spread slowly from the needle tip, this indicated the successful introduction of the needle into the myenteric layer. Twenty to thirty sites were injected in the proximal and distal gastric antrum. An attempt was made to inject the dorsal and ventral aspects of the stomach equally, but injections favored the ventral aspect of the stomach, as the dorsal was less accessible. After completing each infusion, the needle was left in place for 5s before removal to ensure full absorption. After removing the loosing suture attached to the stomach, sterile suture was then applied to the skin.

### Surgical details

The following provides details on viral and drug injections, catheterizations, denervation, and brain electrode implantation. All surgeries were performed in a Biosafety Level 2-approved laboratory. All mice including surgical mice were housed at the Mount Sinai specific-pathogen-free (SPF) Animal Facility. The surgical mice were monitored daily for body weight and food intake. In all cases, preoperative analgesia: 0.05 mg/Kg Buprenorphine (s.c.); anesthesia induced by 3% isoflurane and maintained by 1.5% ∼2% isoflurane; postoperative analgesia: 0.05mg/Kg Buprenorphine (s.c.) twice a day for three consecutive days. The surgical areas were shaved and cleaned with iodine soap and wiped with 70% isopropyl alcohol. All incisions were thoroughly disinfected with a layer of Baytril ointment. All surgeries were performed under stereomicroscopes, with animals placed on a heated pad (CMA 450; Harvard Apparatus, Holliston, MA). After surgery, animals were allowed to recover under infrared heat until they chose to reside in the unheated side of the cage.

### Subdiaphragmatic vagotomy

Animals were 8-hour food restricted before the surgery. The abdomen was shaved and cleaned. A midline incision was made into the abdomen. The liver and stomach were retracted aside to expose the esophagus. The branches of the vagus nerves innervating the stomach were carefully separated from both the esophagus and the left gastric artery, and bilaterally severed with an electrical cauterizer. Sterile suture was then applied to muscle and skin.

### Measurement of gastric pH

To measure gastric pH, a chronic gastric fistula surgical method was adopted from rats. This method allowed repeated observation within a single individual, facilitating measurement of changes in gastric pH that are known to occur with anticipation of food. The abdominal area was shaved of fur, and cleaned with providone iodine and 70% isopropyl alcohol. The stomach was exposed through a midline incision, and bathed in saline with 100 micromolar nicardipine, to reduce its movement and facilitate placement of the stomach drain. A purse-string suture was created at the border of the gastric corpus and fundus, and then a 3 mm incision was made in the center of the purse-string. Residual gastric material was flushed out of the stomach with saline, then the stomach drain was introduced through the purse string, and the string was tightened around the drain, and knotted. Next, the abdominal muscle was sutured around the drain, and then the exterior end of the drain was passed through the skin. The exterior skin was sutured around the drain, trapping a small skirt of surgical polyester mesh which had been previously attached to the drain. The drain was capped with a truncated female leur fitting, which had been sealed with epoxy. After 3 to 4 days of recovery, gastric pH production appeared to recover, with resting pH in ad lib fed animals around 4-5 pH. To measure pH, animals were briefly anesthetized with iso-flurane, the cap sealing the stomach drain was quickly removed, and 600 microliters of saline was flushed into the stomach, and then as much fluid and gastric material as possible was collected by aspiration, the stomach drain was recapped and the animal revived. The retrieved material was mixed first, then briefly spun, 1 min, 15kg rcf, to separate detritus from the gastric secretions. The pH of the separated fluid was measured using Argenthal pH electrode (Mettler-Toledo seven compact pH amplifier connected to Sensor InLab Expert Go probe.)

### Stereotaxic viral injection

Animals at the time of surgery were 6-week-old. Injections were performed with borosilicate glass capillary class pulled into fine needles on a Flaming-Brown Puller (P-97 Sutter). Most injections were performed by manual application of pressure, using filament-free 1.0 mm OD, 0.5 mm ID capillary glass (Sutter B100-50-10). Pipettes were broken back after pulling using a ceramic tile. The pipette openings ranged from 10-20 micrometers. After pulling needles, graduation markers were added to the glass tubing at 1 mm increments, which corresponds to volume graduations of about 220 nanoliters. Needles were connected to polyurethane tubing (Braintree Micro-Renathane® .080″ x .038″) coupled to a 3 or 5 CC syringe, and front filled first with 0.5 microliters of Flurinert FC-40, and then front filled with virus solution. After moving the pipette to the desired stereotaxic coordinate, virus solution was slowly injected by carefully applying pressure to the coupled syringe and visually monitoring the movement of the fluid meniscus under high magnification of a surgical stereo microscope. The volume equivalent of one graduation was voided over the course of 10 minutes, and the pipette was withdrawn 5 minutes after completing the injection. Needles were refilled between sides when performing bilateral injections. Some injections were performed using injection capillary (Drummond 3-000-203-G/X) filled with mineral oil connected to a Drummond nanoject III. Nanoject injections were performed using a program: 5 nanoliters were injected at a rate of 1 nanoliter / second, followed by a 15 second pause. This was repeated 40 times, for an equivalent injection of 200 nanoliters at a rate of 20 nanoliter / minute. Injection placements were verified histologically from sections cut on a sliding freezing microtome. Injections using either method appeared to be qualitatively similar in extent and spatial spread.

### Real-time online place preference

Ethovision 12XT was used to control activation of a DPSLL laser in real-time based on the location of the experimental subject mouse in a divided arena (An 900 cm2 rat cage). Two different textures for the cage floor indicated the two sides of the arena, on one half, a corncob bedding was spread and a vented cage top placed on top of the milled corncob material. The other half of the cage floor was covered with a lab bench top protector (chunx). Animals were acclimated to the chamber for 15 minutes. Most experimental subjects did not express a strong preference between these two floor coverings. After 10 minutes, the less preferred side was paired with 1 Hz laser activation for the ChR2 group (15 millisecond pulse width) and the more preferred side was paired with laser illumination for the GtACR2 group (constant illumination 3-5 mW). After assigning the laser paired zone, the baseline location preference was measured for 5 minutes, before activating the laser. Following this, 10 minutes of laser ‘online’ preference was measured, during which time the laser was activated when the mouse was detected with in the previously selected half of the arena. A mini-io usb output box (EthoVision) was used to trigger activation of the laser, it was configured hold a TTL signal high when the animal was detected in the laser paired zone. Laser pulses were controlled by programming the TDT RZ5 to generate pulses trains gated by the EthoVision TTL. Experiments were performed in a dim room with 850 nM IR illumination and a basler aCA1300 gm (IR cut removed) provided video input for the EthoVision software. Implanted optic fibers (Doric) were connected to a DPSSL 473 nm laser via a fiber optic rotary joint commutator (Doric).

### Open field analysis

A 21 inch diameter circular arena fabricated out of white PVC plastic was illuminated by red led light (630 nm). Animals were not acclimated to the environment, open field behavior was assessed in the novel environment. For each session, animals were placed in the arena for 30 minutes. In the first 20 minutes no laser activation occurred. Next, for 10 minutes laser activation occurred periodically, laser activation lasting 10 seconds repeated every 30 seconds (start to start). After 20 cycles of laser activation, the laser activation ceased, and the animals position was tracked for another 5 minutes. The frequency of laser stimulation for the ChR2 expressing group was set at 10 Hz, pulse duration 15 milliseconds. For the GtACR2 expressing group, laser activation was continuous (2–4 mW). Implanted optic fibers (Doric) were connected to a DPSSL 473 nm laser via a fiber optic rotary joint commutator (Doric).

**Table 1.**
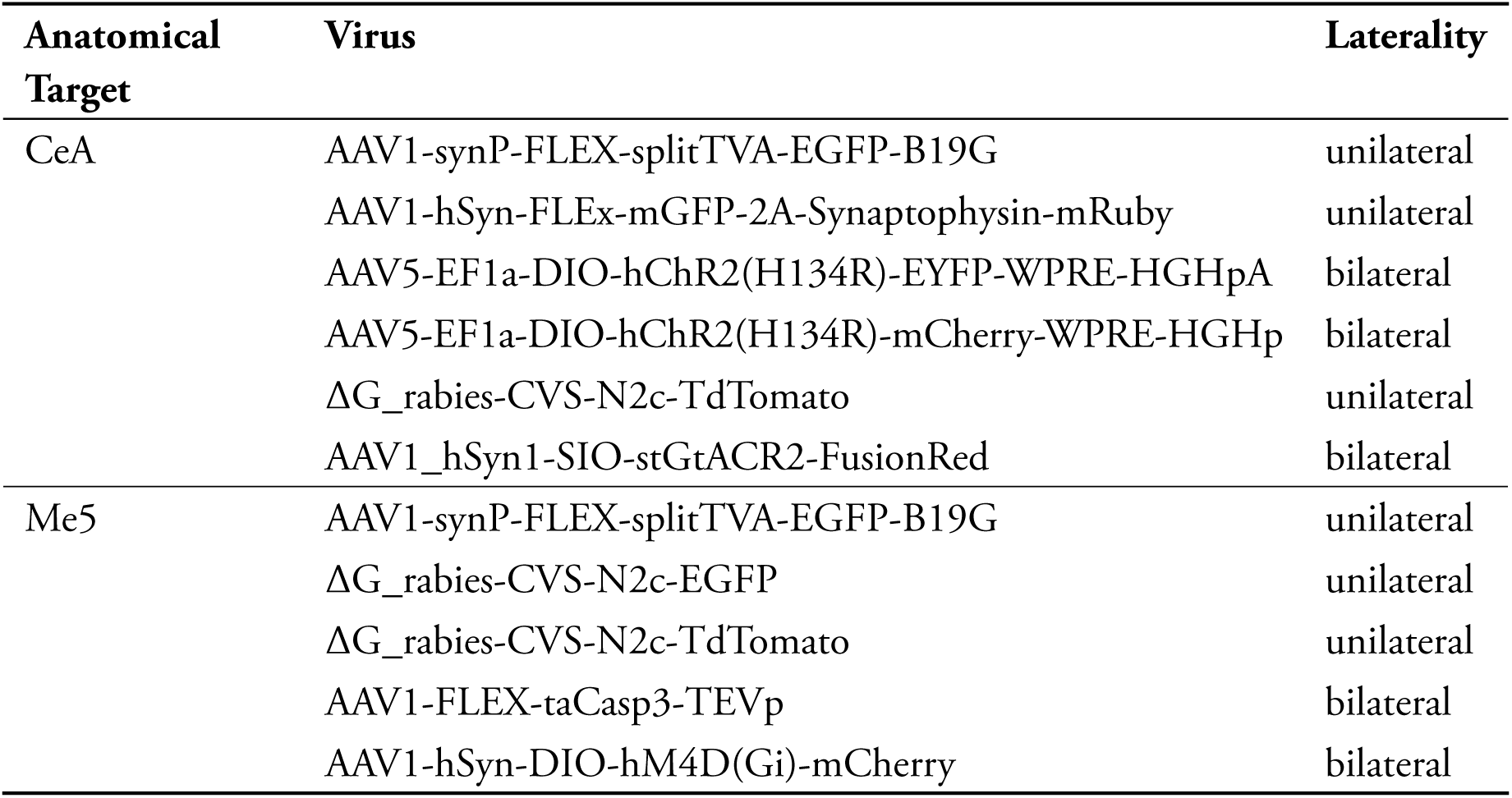
Virus injection table.

**Table 2.** Stereotaxic coordinates used for injections.

| Structure | AP | ML | DV |
| --- | --- | --- | --- |
| CeA | -1.0 | ±2.5 | -4.8-5.2 |
| Me5 | -5.4 | ±1.0 | -3.6-4.0 |
| NTS | -7.5 | ±0.3 | -5.3-5.5 |
| PSTn | -2.4 | ±1.1 | -4.9-5.1 |

### Nose­poking for laser activation

Nose poking was measured in a Med-Associates cages. Nose pokes were measured with Med-Associates IR beam break poke ports (ENV-375A-NPP) detected pokes were recorded, and for pokes on the laser-paired port, the laser was activated in a 10 Hz train lasting 0.5 seconds (15 millisecond pulse duration). Following each poke causing a train of laser stimulation, a 0.5 second inactive period ensued, during which pokes were recorded but they would not trigger another train. After this 0.5 second inactive period the next poke on the paired port would trigger another train. Pokes during the inactive period did not cause a time out. Implanted optic fibers (Doric) were connected to a DPSSL 473 nm laser via a fiber optic rotary joint commutator (Doric).

---

**Figure S1:**
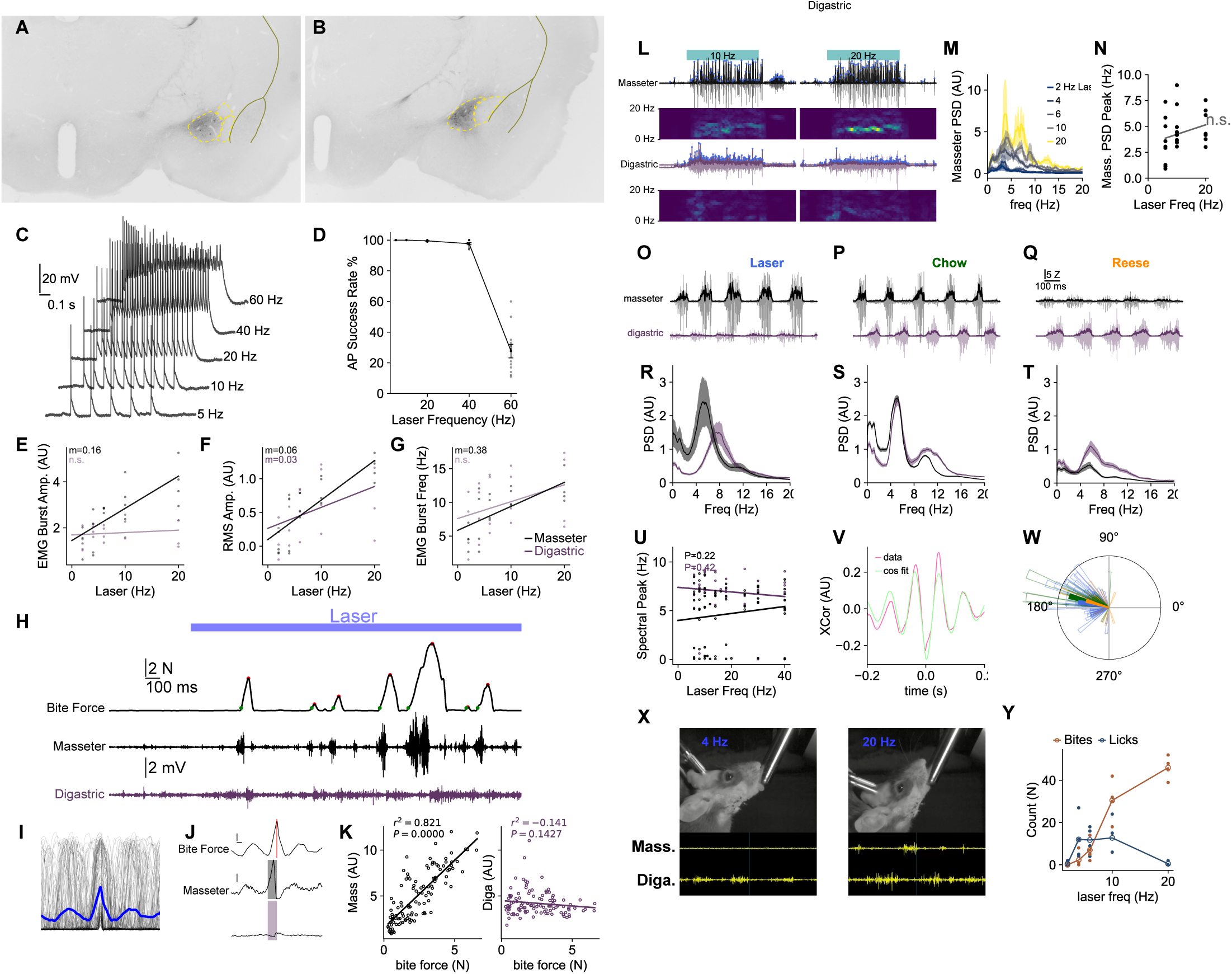
Stimulation of CeA^Isl1+^ neurons drives rhythmic contraction of the jaw closer masseter muscle. (A-B) Representative images of GFP expression following injection of Cre dependent AAV into the central amygdala of Isl1^Cre^ animals. Note that viral expression is restricted to the medial division of the central amygdala. (C) Representative traces from whole cell patch clamp recordings of CeA neuron expressing eYFP following injection of AAV driving Cre dependent expression of ChR2.eYFP in Isl1^Cre^ mouse. Response to trains of 470 nm LED flashes. (D) Quantification of fraction of flashes within a train successfully evoking action potentials (N=12 cells, 3 mice). LED stimulation at 60 Hz lead to depolarization blockade and failure to repetitively fire. (E-G) Quantification of jaw opener (digastric) and closer (masseter) EMG signal in response to stimulation of CeA^Isl1+^ neurons. (E) Average amplitude of each burst peak is linearly related to the frequency of CeA^Isl1+^ neuron stimulation for the masseter muscle, (N=4, P=1.41e-08, slope=0.156667), but this relationship is not significant for the digastric muscle, (N=4, P=0.528). (F) In contrast, the averaged total area under the rectified EMG (RMS amplitude) is linearly related to the frequency of CeA^Isl1+^ neuron stimulation for both masseter muscle, (N=4, P=8.169-10, slope=0.062534), and for the digastric muscle, (N=4, P=0.028, slope=0.031494). (G) The average frequency of EMG bursts is linearly related to the frequency of CeA^Isl1+^ neuron stimulation for the masseter muscle, (N=4, P=2.212e-06, slope= 0.384417), but not the digastric muscle, (N=4, P=0.115). (H-K) The masseter EMG signal is linearly related to bite force. (H) Example traces of signals recorded during activation of CeA^Isl1+^ neurons. Upper trace, bite force, middle trace, masseter EMG, lower trace, digastric EMG. Blue bar above indicates initiation of laser stimulation. Red dots in the upper force trace show detected peaks in the force recording, which are used for later alignment of the EMG and force data. Green dots in the force trace indicate the time of threshold crossing for the alternate method of alignment. (I) Each peak in the force trace (red dots in A) is aligned to time zero, individual traces are overlayed in black, and the average force is plotted in blue. (J) Triggered averages of individual bites are computed for bite force (top), masseter EMG (middle), and digastric EMG (bottom). Individual bites are aligned so that the peak of recorded force is at time zero, shown by the red line. Orange patch overlaying the masseter EMG trace and blue patch overlaying the digastric EMG trace show the period of during which the EMG signal was summed for the purpose of creating the force - EMG correlation. (K) The masseter EMG activity has a tight positive correlation with measured bite force (left plot), whereas digastric EMG activity is not significantly correlated (right plot). (L-N) Rhythmic masseter EMG activity evoked by stimulation of CeA^Isl1+^ neurons has a peak in spectral power that is insensitive to the rate of laser stimulation. (L) Example traces showing activation of the masseter (black trace top), and the digastric (purple trace) in response to a increasing frequencies of CeA^Isl1+^ neuron stimulation. Rectified integrated EMG signals (thicker lines) are used for the spectral analysis. Upper (masseter) and lower (digastric) spectrograms show the emergence of regular rhythmic activity between 4 and 10 Hz during stimulation that is marked for the masseter during stimulation of CeA^Isl1+^ neurons between 10 and 20 Hz. (M) Power spectral density estimates for masseter during stimulation of CeA^Isl1+^ cells at increasing stimulation rates. (N) No significant linear relationship exists between the predominant frequency peak of masseter spectral activity and laser stimulation rates. (O-U) Stimulation of CeA^Isl1+^ neurons releases rhythmic activity of the masseter muscle that has similar spectral characteristics as is observed in the consumption of chow. (O-Q) Representative EMG recordings during activation of CeA^Isl1+^ neurons by laser, during chow consumption, and during reese’s peanut cup consumption. (R-T) Aggregate data (N=5) of masseter and digastric spectrograms corresponding to panels O,P,and Q respectively. (U) As in the previous group (Panel N), no significant different is observed between the frequency of predominant peak in spectral activity and the rate of laser stimulation, for either the masseter or digastric muscle. (V-W) The phase relationship between the masseter and digastric muscle during stimulation of CeA^Isl1+^ neurons is similar to that observed during consumption of food. V. Example data showing representative fitting of a damped cosine function to the cross-correlogram of the masseter and digastric recording. W. Radial plot of fit phase term (plot angle), and amplitude term (bar height). (X-Y) Stimulation of CeA^Isl1+^ neurons sometimes released licking behaviors. X. Example still frames from videos recorded while mouse was presented with dry sipper and the laser was activated at 4 (left frame) or 20 Hz (right frame). In this mouse, EMG from the masseter and digastric were recorded simultaneously. A lick is elicited with 4 Hz stimulation, whereas at 20 Hz the mouse is biting the sipper (note the masseter activation). (Y) Quantification of number of bites and licks evoked by different rates of laser activation ranging from 2-20 Hz, when the mouse is presented with a dry sipper.

**Figure S2:**
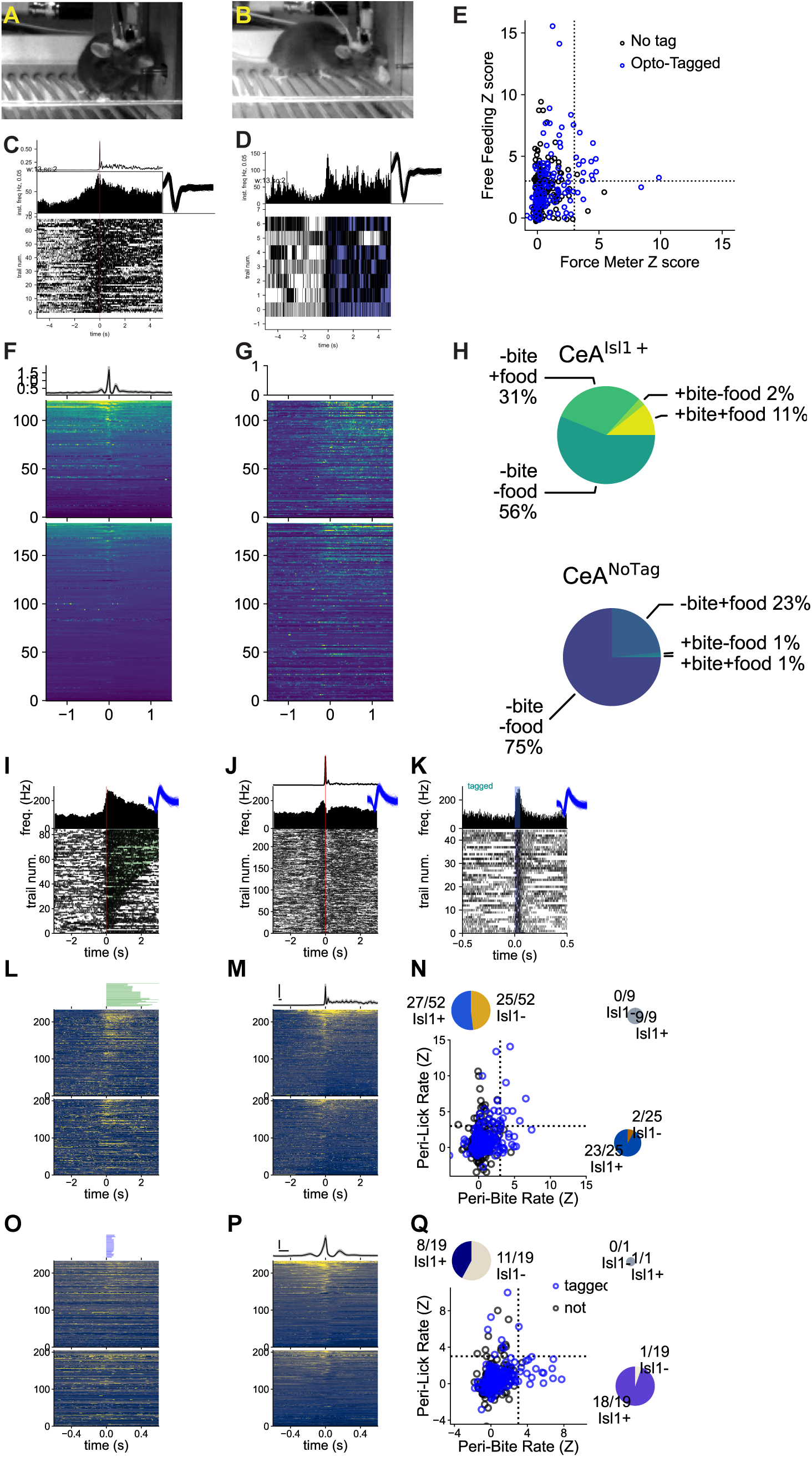
Opto-tagged CeA Isl1+ units are active during forceful biting, and during biting of chow. A-B. In some recording sessions, following recording of unit activity during operant biting of force meter, chow was added to the cage and neurons were recorded during consumption of chow. Video analysis of these sessions was used to assign onset of feeding times. Here a frame from the video is shown during the first segment of the session, where the animals is biting the force meter for an infusion reward. B. Frame from video marked as time of biting food comparable to A. C. Raster showing activity of a representative opto-tagged unit during biting of the force meter. D. Raster showing activity in response to initiation of a feeding bout. Same unit as shown in (C). E. Comparison of Z transformed firing rates during biting of the force meter (X axis) as compared to biting of chow (Y axis) for the each unit recorded with data for both behaviors. Dashed lines show an significance cut off at 3 Z units. Note that opto-tagged units are active both during consumption of food and forceful biting of the strain gauge. F. Heat map of aggregate activity of all units during biting of force meter. color scale −1 : 3 Z units. Top heat map is from opto-tagged cells, bottom heatmap is from the non-tagged cells. Trace above the heat map shows the average force applied to the force meter for all aligned bite events. G. Heat map of aggregate activity of all in responses to bouts of chow consumption. As in (F), top heat map is opto-tagged cells, bottom heatmap is non-tagged cells. No force is applied to the force meter here. Row order of the units is the same as in the heatmaps in (F). H. Pie plot showing the relative number of cells that are significantly active during biting of the force meter, and during biting of food, for both opto-tagged cells, and non-tagged cells. I,L,N. A subset CeA^Isl1+^ neurons are active during bouts of licking. I: Raster plot of representative opto-tagged unit aligned to onset of bouts of licking the water sipper. Green boxes overlayed on raster indicate the onset and offset of each licking bout. L: Heatmap of Z transformed firing rates for opto-tagged (upper) and non-tagged (lower) units, aligned to lick bout onset. Rows of heat map are ordered according to row order in panel M. Upper green boxes show the lick bouts for each record. Box height is proportional to number of bouts, box width is proportional to averaged duration of bouts. (M) Heatmaps of Z transformed firing rates for tagged (upper) and non-tagged (lower) units, aligned to ‘bouts’ of biting. Rows are ordered by the first principle component computed on the firing rate responses during bite bouts. N. Comparison of Z transformed firing rates during bouts of biting the force meter (X axis) as compared to bouts licking the water sipper (Y axis) for the each unit recorded with data for both behaviors. Dashed lines show the significance cut off at 3 Z units. Note that relatively fewer opto-tagged units are active during both bouts of licking and bouts of biting of the strain gauge. O-Q. Units that are significantly active during individual bites, do not show significant rate modulations about the emission of individual licks. O: Heatmap of Z transformed firing rates for opto-tagged (upper) and non-tagged (lower) units, aligned to individual licks. Rows of heat map are ordered according to row order in panel P. Upper blue boxes show the averaged lick contact time for each record. Box height is proportional to number of licks, box width is proportional to averaged duration of contact to the lick sipper. Q. Comparison of Z transformed firing rates during bites on the force meter (X axis) as compared to licks on the water sipper (Y axis) for the each unit recorded with data for both behaviors. Dashed lines show the significance cut off at 3 Z units. Note only a single unit is significantly active during both behaviors.

**Figure S3:**
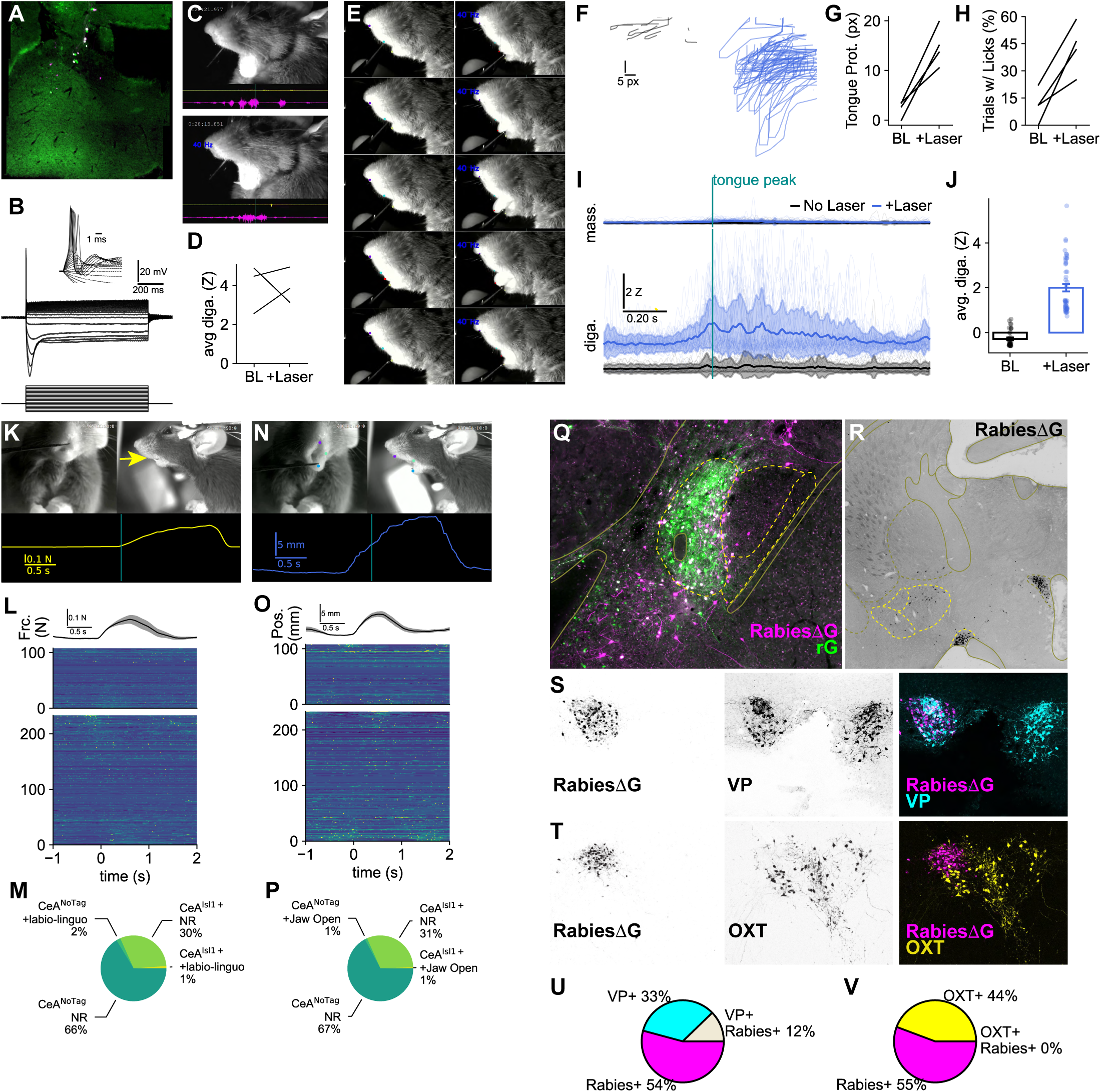
CeA::Is1l+ neurons do not respond to orofacial stimulation, and have a pattern of synaptic input suggesting limited direct sensory input. A. Example micrograph of Me5 rabies starter neurons, cells expressing the rG helper (green) and rabies (magenta) appear white. These neurons are the starter cell population for mono-synaptic retrograde tracing that identified CeA^Isl1+^ cells as presynaptic. B. Example recording of family of current steps used to identify Me5 neuron by intrinsic electrophysiological properties. Note the prominent sag with hyperpolarizing steps, and the short time scale membrane resonance. C-D. Stimulation of CeA^Isl1+^ neurons does not suppress the Jaw Opening Reflex (JOR). C. Representative frame showing the JOR evoked by pressing on the hard pallet. Upper panel no stimulation, lower panel JOR evoked during 40 Hz stimulation of CeA^Isl1+^ neurons. D. Quantification of total digastric (jaw opening muscle) EMG signal elicited by pressing on hard pallet. E-I. Stimulation of CeA^Isl1+^ neurons in urethane anesthetized mouse can potentiate appetitive licking in response to brushing the upper lips. E. Two sequences of still frames showing potentiation of licking by laser stimulation during lip stimulation. Left columns of still frames shows responses to lip brushing without laser activation. Right columns shows responses to a similar manual stimulation of the upper lips which occur simultaneous to high frequency activation of CeA^Isl1+^ neurons by laser activation. Tongue position was tracked by DLC, small red dot on tongue tip shows location assignments. F. Comparison of tongue trajectories elicited by upper lip stimulation without (black) and during (blue) laser activation of CeA^Isl1+^ neurons. Tongue protrusion distance is increased by laser activation (G) as is likelihood to elicit licking (H). I. Contraction of the digastric muscle (as measured by EMG) in response to lip touching is increased by concurrent CeA^Isl1+^ activation. Trials are aligned by the timing of the peak of tongue protrusion. Digastric EMG is significantly greater when laser stimulus is added to the lip brushing stimulation (J). K. Video visualization of experimental set up to measure single unit activity in the central amygdala during stimulation of the teeth. Example of stimulation of the incisor teeth in the labio-linguo direction. Upper frames show synchronized head-on and side view video recording. The yellow arrow on the right video frame indicates the direction of force applied to the incisors. The probe was pressed against the labio surface of the teeth. Lower yellow trace shows the force applied by manual stimulation to the incisor, upward deflection indicates the labio-linguo direction. The vertical cyan bar transecting the force trace indicates the time of the above video frames. N. Example of trial where the lower jaw is manually opened, by gently depressing the mandible. Upper frames show synchronized video record, note the purple, green, and blue dots on the video frames, these indicate the position of tracked features on each frame of video. The blue trace below the video frames plots the displacement in the position of the blue dot, from its resting position. L. Summary heat map data showing single unit responses to stimulation of incisor teeth in the labio-linguo direction. Black trace above the heat maps shows the average force applied across each recording site, gray shading indicates standard error of the mean. 341 cells in total were recorded from three mice. Four site were recorded in each mouse, for a total of 12 distinct recording locations. Of the recorded cells, 108 were classified as Opto-tagged by the SALT criteria, they are labeled as CeA::Isl1+. The remaining 233 cells are labeled as CeA::NoTag. The upper heat map shows responses of the CeA::Isl1+ cells to the tooth stimulation, the lower heat map shows the CeA::NoTag cells. The heat map spans from +3Z (yellow) to −1.5 Z (dark blue). O. Summary heat map data showing single unit responses to manual opening of the jaw. Black trace above the heatmaps shows the movement of the mandible from its rest position, averaged across the recording sites. Upper heat map shows responses of CeA::Isl1+ (opto-tagged cells), lower heat map shows responses of CeA::NoTag cells. M. Pie plots showing proportion of CeA::Isl1+ and CeA::NoTag that significantly respond to stimulation of incisors in the labio-linguo direction. P. Pie plots showing proportion of cells significantly responding to manual jaw opening. Q-V. Rabies virus mapping of inputs to CeA^Isl1+^ cells Q. Example of starter population for synaptic input mapping in the central medial amygdala. Rabies G protein is expressed by Cre dependent recombination in Isl1^Cre^ mice. A small injection of the rabies G helper virus labeled the medial division of the central amygdala. Subsequent injection of ΔGrabies expressing RFP (shown in magenta), double labeled cells (white) are starter population for which synaptic inputs are mapped. 4 mice were included, all cases where similar in the extent and location of the starter injection, all injections are unilateral. R. Tiling image of the anterior portion of the central amygdala, showing the prominent inputs from the Supraoptic Nucleus (SON), and the Periventricular Hypothalamus (PVH). S-V. PVH neurons projecting to CeA^Isl1+^ neurons comprise vasopressin expressing neurons but not oxytocin expressing neurons. S. Representative immunostaining for vasopressin in PVH sections showing rabies virus retro-synaptic labeling from CeA^Isl1+^ neurons. T. Representative immunostaining for oxytocin in rabies injections cases. U. Quantification of PVH cells positive for rabies, for vasopressin, or both. V. Quantification of PVH cells positive for rabies, for oxytocin, or both.

**Figure S4:**
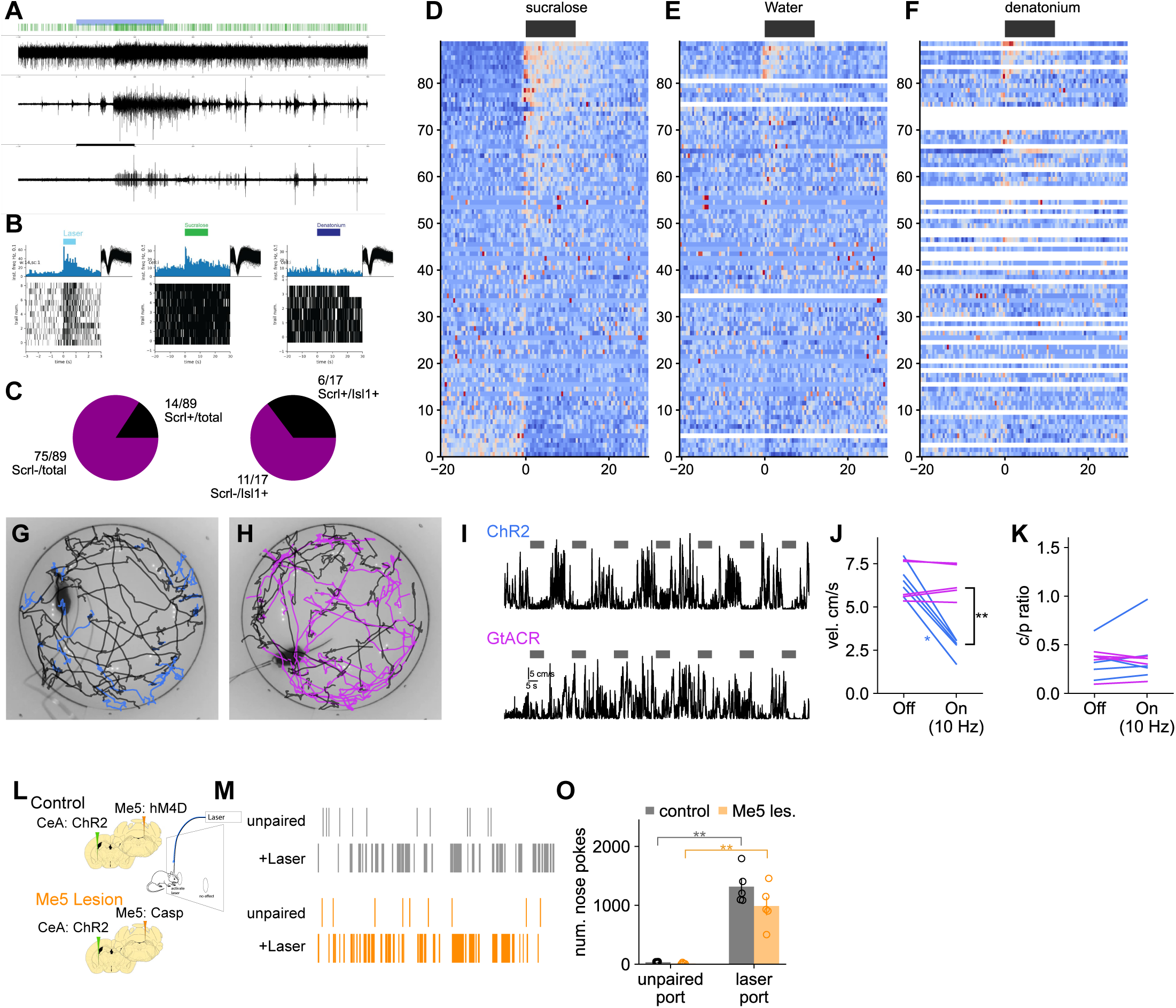
Central Amygdala neurons expressing Isl1 neurons are activated during consummatory motor responses evoked by sweet tastes. Activation of CeA:Isl1+ neurons halts ongoing locomotion in an open field. A. Example trace of single unit recording of optically tagged putative CeA^Isl1+^ unit. Top trace is single unit recording, green ticks above top trace are spikes for an identified unit, the cyan bar indicates the infusion of sucralose into the oral cavity. Middle trace is digastric EMG, bottom trace is masseter EMG. The burst of EMG activity indicates the initiation of an appetitive oral response. B. Raster of responses to laser stimulation (left), sucralose (middle) and denatonium (right). Rasters to taste responses are aligned to the initiation of EMG activity observed after infusion of the tastant. C. Proportion of cells responding to sweet tastes is higher among optically tagged CeA^Isl1+^ than in the general population of amygdala neurons, fishers exact test [P=0.024]. D-F. Heatmap of all recorded neurons responses to sweet (D), water (E), and the bitter tastant, denatonium (F). G-K. Stimulation of CeA^Isl1+^ neurons reduces ongoing locomotion in an open field arena. G. Representative position tracks over 8 trials of laser stimulation in the open field. Each trail is represented by a single track, the first 8 seconds of the track, during which the laser is off are colored black, the subsequent 8 seconds of the trail, during which the laser is on, is colored in royal blue (ChR2 group). Laser activation at 10 Hz essentially dramatically reduces ongoing locomotion. H. Similar to G, but showing representative tracks for 8 trails of laser activation for the animals expressing GtACR in CeA:Isl cells. The portion of the trial track occurring during laser activation is colored in magenta (GtACR group). The laser in continuously on rather than pulsed at 10 Hz in this group. I. Upper trace: Representative velocity traces over time during 7 trails of laser activation for an animal expressing ChR2 in the CeA:Isl1 cells. Period of 10 Hz laser activation are shown by the presence of gray bar above the velocity trace, shared scale bar in lower trace. Lower trace: similar to upper, but animal is expressing GtACR in CeA:Isl1+ neurons, and laser is continuously active during periods shown by gray bar. J. Higher frequency activation (10 Hz) of CeA^Isl1+^ neurons in an open field arena significantly decreased velocity, but inhibition had no effect on walking speed F_(1,8)_=177.22, P=9.7*10^-7, interaction OPSIN X LASER. Pairwise tests showed that the significant difference in walking speed between the two group (activation and inhibition) was only observed during the laser active period (t_(8)_=-7.3,P=0.0000837), during the laser off periods, the two groups walking speeds were not significantly different (t_(8)_=0.29,P=0.78) K. Higher frequency CeA activation (10 Hz) or inhibition (constant illumination 3-5 mW) did not significantly shift animals localization within the open-field arena between the center and the periphery. (mixed ANOVA F_(1,8)_= 1.63, P=0.237 interaction OPSIN X LASER) O-Q. CeA:Isl1+ neuron activation continues to act as an operant reinforcer in animals that cannot generate high force bites. O. The operant self-stimulation paradigm is the same as in panel L, but the viral expression and surgical approach has been extended: in addition to expression of ChR2 in CeA:Isl1+ cells, a Cre dependent caspace virus was also injected in the mesencephalic trigeminal nucleus, (Me5). Because Me5 neurons also express Isl1, they will contain Cre, and the Caspase virus will lesion these neurons. P. Example of poke events into the laser paired port and unpaired port, Similar to M, except the upper event plot (gray) is from a representative Control animal, expressing HM4:mCherry in Me5 neurons. Lower event plot (orange) is from a representative animal expressing with Me5 neurons lesioned via Caspase expression. Q. Control and Me5 lesioned animals nose-poked into the laser paired port significantly more than the unpaired port, mixed ANOVA: pokes ∼ port * lesion + Error(subject/port), port: F_(1,8)_=116.09. No significant interaction was observed between the lesion and port: F_(1,8)_=2.14,P=0.182. Post hoc tests confirmed that both animals in the control (t_(4)_=-9.27,P=0.00075) and intact (t_(4)_=-6.196,P=0.0034) groups poked more into the laser paired port than the unpaired port. Further, post-hoc tests comparing the pokes into the laser paired port between the lesion and control groups did not reveal a significant difference (t_(8)_=1.601,P=0.1479).

**Figure S5:**
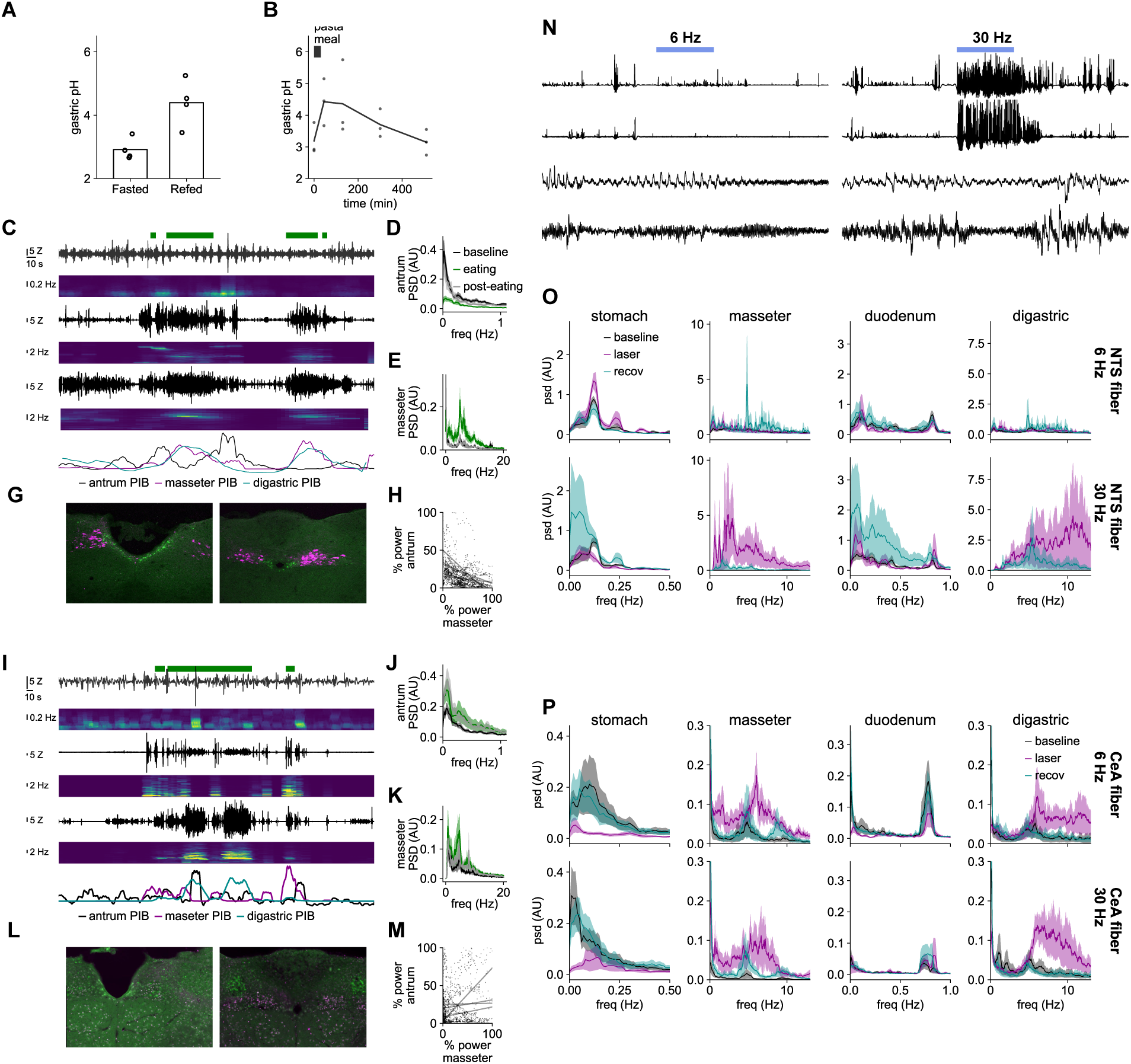
Suppression of the gastric antral rhythm observed during CeA^Isl1+^ stimulation is also observed during consumption of food, this phenomenon depends on the integrity of the vagus nerve. A-B. Gastric pH is increased by the consumption of food. A. Gastric pH was measured by terminal collection of gastric contents either after a 12 hour fast, or 2 hours after last access to food. Faster gastric pH is lower than the re-fed condition. B. Measurement of time course of increase in gastric pH following a pasta meal. Chronic gastric fistula animals (N=3). B. Representative EMG recordings of gastric and jaw signals before, and during, consumption of standard chow. Traces and spectrograms are in pairs: upper is gastric antrum, middle is masseter, lower is digastric. Note the frequency range for the gastric antrum spectrogram is much lower than for the masseter and the digastric. Green bars above top trace indicate the period of time animal is consuming chow. Line plots at the bottom show the total power in the relevant frequency band for the gastric antrum and for the maseter. D. Consumption of food suppresses the electrical rhythm of the gastric antrum, the amplitude of the power spectrum between 0 and 0.3 Hz is significantly reduced (t[4]=7.06, P=0.0021) during food consumption as compared to the baseline period immediately preceding consumption. E. Power spectral changes in masseter activity during food consumption. F. A significant negative correlation between the amplitude of the power spectrum of the antral rhythm and the amplitude of the chewing rhythm in the masseter is observed in 5/5 intact animals. G. Retrograde transport of cholera toxin subunit b (ctb) following injection in the stomach in an intact animal. I-M. Gastric motility is not suppressed during chow consumption in animals receiving a bilateral subdiaphragmatic vagotomy. I. Example traces as in panel B. J. Power spectrum as in panel D, note that while food consumption has an effect in vagotomized animals, (F(2,y)=6.9, P=0.028), post hoc tests suggest an increase in gastric antral activity immediately following consumption (t_(3)_=-4.3, P = 0.023). K. Power spectrum as in C, in vagotomized animals. M. None of vagotomized animals had a significant negative correlation between masseter and antral activity. L. Following ctb injection in the stomach of vagotomized animals, very little labeling is observed in the dorsal vagal complex. N. Representative traces EMG activity of the (from top to bottom): digastric, masseter, gastric antrum, and duodenum in response to stimulation of CeA^Isl1+^ terminal fibers in the NTS at 6 Hz (left) and 30 Hz (right). Note the potentiation of the gastric rhythm in response to 6 Hz, as compared to the inhibition observed in response to 30 Hz stimulation. O. Average power spectral density estimates from mice with fiber position in above the NTS for stimulation of CeA^Isl1+^ terminals projecting to this region. Upper row, are changes in response to laser stimulation at 6 Hz, lower row are changes in response to 30 Hz. P. Average power spectral density estimates from mice with fiber positioned above the central amygdala for stimulation CeA^Isl1+^ cells independent of projection target, plots are otherwise comparable to those in panel O, save absolute amplitudes are different.

